# Enhancing timber traceability via multielement and strontium isotope ratio: An example from the Eastern Alps

**DOI:** 10.1101/2024.09.13.612829

**Authors:** Agnese Aguzzoni, Francesco Giammarchi, Ignacio A. Mundo, Giulio Voto, Giustino Tonon, Werner Tirler, Enrico Tomelleri

## Abstract

International timber trading is subject to rigorous certification schemes that require the disclosure of essential information, including the tree species and geographic origin of the timber in question. Regrettably, the lack of readily accessible forensic tools to verify compliance has facilitated the proliferation of illegal timber trading, with dramatic consequences for ecosystems and biodiversity. The objective of this study was to investigate the potential of a multichemical approach based on the multielement and strontium isotope (^87^Sr/^86^Sr) ratio analysis combined with chemometrics to test sample recognition according to their species and geographic origin. The sampling area covered a portion of the Eastern Alpine region, which is characterised by a significant economic reliance on wood. The study focused on three representative species from local forests: Norway spruce, European larch, and Swiss stone pine. Samples were characterised from stands grown on diverse bedrock types. Our findings revealed a strikingly consistent variation in the multielement profiles across different species, thereby enabling flawless sample recognition. Considering the geographic origin, the ^87^Sr/^86^Sr ratio proved to be a pivotal parameter, by virtue of its correlation with the geo-lithological composition of the growing area. Combining the chemical markers, an accurate sample classification based on multiple decision trees was attained, even comparing forest stands grown on the same bedrock type. These findings offer novel insights into the utilisation of chemical markers in provenancing and authenticity studies, thereby enhancing the adoption of integrated approaches to counteract illegal timber trade.

## 1. Introduction

In the context of global change, the demand for renewable resources is continuously increasing (Fekete et al., 2021; Leirpoll et al., 2021). Wood can be successfully employed as a substitute for fossil fuels in the production of energy, as well as for materials with a higher carbon footprint in the construction industry (Gustavsson et al., 2021). On the other hand, elevated wood harvesting rates necessitate the implementation of sustainable forest management strategies. This is to prevent deforestation and promote the availability of forestry-related products in compliance with legal requirements (Fekete et al., 2021; Lowe et al., 2016). In this regard, forest certification schemes for sustainable management and chain-of-custody have been demonstrated to enhance the prevention of illegal activities (Paluš et al., 2018). Moreover, governments have established measures to address the issue of indiscriminate or illegal logging and trade. Notable examples include the Convention on International Trade in Endangered Species of Wild Fauna and Flora (CITES) and, more recently, the European Union Forest Law Enforcement, Governance and Trade (EU-FLEGT) Action Plan. Nevertheless, these are still critical issues in the timber market (Moral-Pajares et al., 2020), with the potential to undermine forest ecosystem conservation, leading to adverse effects on the ecological status of forests as well as on the economic and social well-being of local communities (Ceccon et al., 2020). This is particularly the case with tropical timbers, with estimates suggesting that approximately 30-90% of these products in the market originate from illegal harvesting (Deklerck et al., 2020; Vlam et al., 2018). In Europe, illegal logging and trading persist as significant challenges, endangering the limited remaining pristine and old-growth forests on the continent (Bouriaud, 2005). Consequently, accurately assessing the exact tree species and its origin is vital for effectively contrasting illegal logging activities in the forestry sector. Furthermore, determining provenance can encourage the use of premium timber in accordance with sustainable forest management systems and from specific geographical regions.

Several techniques have been developed to detect the species and verify the provenance of traded timber, with a focus on its inherent characteristics. These include, for instance, DNA analysis, mass spectrometry, and isotope analyses (Dormontt et al., 2015). All these methods have both advantages and limitations. The use of DNA analysis allows for the identification of a sample at the genus or species level, as well as the geographical area of origin at a fine scale. However, this may only apply to natural populations with a sufficient degree of genetic diversity, and may not be applicable to plantations with exogenous genetic material (Kagawa and Leavitt, 2010). Furthermore, it necessitates the availability of extensive reference datasets (Deguilloux et al., 2003) and considerable laboratory work. The use of mass spectrometry, specifically Direct Analysis in Real Time Time-of-Flight Mass Spectrometry (DART-TOFMS), has recently demonstrated efficacy in the rapid identification of species, but its potential for provenancing remains uncertain (Deklerck et al., 2020; Evans et al., 2017; Paredes-Villanueva et al., 2018; Price et al., 2022). Other mass spectrometry techniques capable of determining the multi-element or multi-isotope ratio fingerprint have proven highly valuable in provenance studies (Balcaen et al., 2010; Benson et al., 2006). The multi-element fingerprint of vegetal samples derives from that of the growing soil and may indeed provide univocal information about the place of origin (Boeschoten et al., 2023b). Similarly, the isotope ratios of several elements have been employed to ascertain the provenance of timber, demonstrating their efficacy (Boeschoten et al., 2023a; Erban Kochergina et al., 2021; Gori et al., 2018). Among the various isotope systems, the analysis of strontium isotopes (^87^Sr/^86^Sr) has been identified as a promising approach for discriminating the origin of timber, particularly at a fine scale (D’Andrea et al., 2023; Vlam et al., 2018). This is because strontium is a soil-derived marker that depends on the local geo-lithology (Hajj et al., 2017). Although this isotope ratio has been extensively employed for the tracing of food products (Baffi and Trincherini, 2016), there is a lack of examples concerning its application in wood provenancing, with most available studies focused primarily on the field of archaeology (English et al., 2001; Erban Kochergina et al., 2021; Million et al., 2018; Reynolds et al., 2005; Rich et al., 2016).

In the Alpine region, the timber supply chain constitutes a significant component of the local economy (Gori et al., 2018). Concurrently, despite the prevalence of a limited number of dominant species, predominantly softwoods such as Norway spruce or European larch, the landscape exhibits considerable diversity. This is attributable to several factors such as altitude, morphology, and geological setting (Ohler et al., 2020). All these drivers influence tree growth, consequently affecting their quality at a fine spatial scale. An illustrative example is the high-quality Norway spruce tonewood, which is found only in a few geographically limited areas in the Alps, where specific growth conditions contribute to the outstanding acoustic properties of this material (Cherubini et al., 2022; Romagnoli et al., 2003). This makes the possibility of detecting and certifying the provenance of locally produced timber appealing. To gain deeper insight into this topic, this study analyses the multielement composition and ^87^Sr/^86^Sr ratio of samples from three important species for the Alpine timber economy: Norway spruce, European larch, and Swiss stone pine, which originate from different locations in the Eastern Alps. The objective of this study is to assess the effectiveness of combining chemical analyses with chemometrics for differentiating between species and origins of the wood samples. We conducted a demonstrative investigation at a limited geographical scale to demonstrate the efficacy of our approach in discriminating neighbouring growth sites with the potential for significant implications for the development of a novel standard multi-chemical certification of origin.

## 2. Materials and Methods

### 2.1 Sites description and sampling

We selected ten different managed forest stands in the Eastern Alps. Specifically, the study area encompasses South Tyrol (Italy) and the neighbouring Italian regions of Lombardy, Trentino, and Veneto, in addition to a stand in Tyrol (Austria) (Fig. 1). The stands located in South Tyrol were characterized by diverse bedrock types, whereas the stands in the other regions were situated within the dolomitic terrain (Table 1). The stands were selected based on the typical Alpine high-mountain mixed conifer forest composition, which included Norway spruce (*Picea abies* Karst., PIAB), European larch (*Larix decidua* Mill., LADE) and Swiss stone pine (*Pinus cembra* L., PICE). The three species were present in varying proportions across the sites, though Norway spruce was consistently the most abundant. The slope of all sampling sites was approximately 50-60%. The altitude of the sampling sites ranged from 1,000 to 2,000 m a.s.l, with variations in slope aspect across the different locations. Particularly, due to the specific requirements of Swiss stone pine, lower altitudes corresponded to colder conditions, with stands typically growing on north-facing slopes (Table 1).

The forest structure exhibited similarities across the sampling sites, displaying a monolayered or bi-layered vertical structure, a random spatial structure, and a regular or dense forest density. In each site, five dominant or co-dominant trees were selected for each species, besides the sampling site n. 4, where only four larches were sampled, and the site n. 6, which lacked Swiss stone pine. The diameter at breast height (DBH, 1.3 m) of trees ranged from 16 to 39 cm, from 17 to 31 cm, and from 18 to 39 cm in European larch, Norway spruce, and Swiss stone pine, respectively. All of the aforementioned measures align with the standard commercial log sizes set by the local regulatory authority (Provincia Autonoma di Bolzano, 2006). A wood core was extracted from each tree at breast height with a 10-mm increment borer (Haglöf, Sweden).

**Table 1.** Characteristics of the sampling sites.

| **Siten.** | **Region** | **Location** | **Sampling**<br>**Year** | **Bedrock**<br>**type** | **Altitude**<br>**(m a.s.l.)** | **Aspect** |
| --- | --- | --- | --- | --- | --- | --- |
| 1 | South Tyrol | Laces | 2020 | Gneiss | 1820 | NW |
| 2 | South Tyrol | Lavazè | 2020 | Porphyry | 1650 | NW |
| 3 | South Tyrol | Passo Nigra | 2020 | Mix | 1930 | SW |
| 4 | South Tyrol | Plose | 2020 | Phyllite | 1630 | N |
| 5 | South Tyrol | Val di Vizze | 2020 | Calc-schists | 1920 | SW |
| 6 | Lombardy | Borno | 2021 | Dolomite | 1060 | E |
| 7 | Lombardy | Livigno | 2021 | Dolomite | 1940 | NW |
| 8 | Trentino | Pampeago | 2021 | Dolomite | 2100 | E |
| 9 | Tyrol | Hahntennjoch | 2021 | Dolomite | 1820 | W |
| 10 | Veneto | Passo Tre Croci | 2021 | Dolomite | 1840 | N |

**Figure 1.**
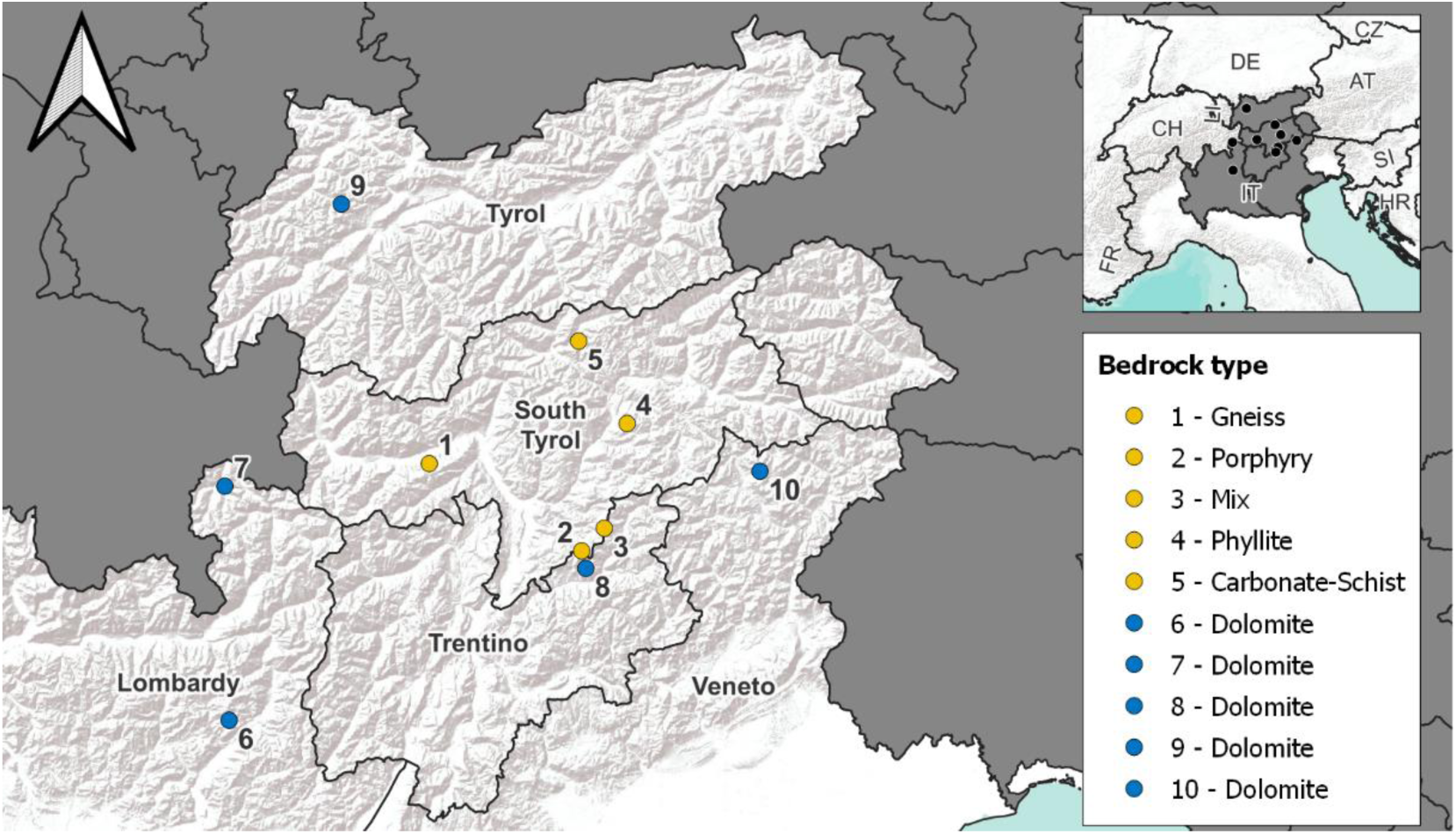
Localization of sampling sites and bedrock type. See Table 1 for more information.

### 2.2 Multi-element analysis

Wood core samples for multi-element and isotope analysis were oven dried (40 °C until constant weight) and, after removing the bark, pulverised and homogenised in a zirconia ball mill. A sample aliquot (0.5 g) was dispersed in 2.5 mL of HNO_3_ (65%), purified by quartz sub-boiling distillation system (DuoPur, Milestone, Sorisole, Italy), and digested in 15 mL PTFE tubes closed with PTFE caps using a microwave digestion apparatus (UltraWave, Milestone, Bergamo, Italy). The digestion cycle consisted of a temperature ramp to 200 °C over 25 min, followed by an isothermal phase of 10 min at a constant power (1500 W). A cleaning cycle of the tubes with HNO_3_ (65%, subboiled) was performed between digestion cycles. The digest solutions were transferred to graduated disposable vials and brought to the final volume with high-purity deionized water (18.2 MΩ cm) (Elix, Millipore, Billerica, MA, USA), which was additionally purified through the DuoPur distillation system. Subsequently, an aliquot of each digest solution was diluted to a final concentration of HNO_3_ of 2% and analysed at the inductively coupled plasma mass spectrometer (iCAP Q ICP-MS) (Thermo Scientific, Bremen, Germany) equipped with an autosampler ASX-520 (Cetac Technologies Inc., Omaha, NE, USA) for multi-element quantification. The multi-element analysis included the following elements: B, Mg, Al, P, K, Ca, Ti, V, Cr, Mn, Fe, Co, Ni, Cu, Zn, Se, Rb, Sr, Mo, Ag, Cd, Sn, Ba, Tl, and Pb. Mono and multi-element standard solutions were used to prepare a seven-point calibration curve in the following ranges: 0.01 – 20 µg/mL for Ca, 0.005 – 10 µg/mL for K and Mg, 1 – 2000 ng/mL for P, and 0.1 – 200 ng/mL for the remaining elements. A solution of Sc, Ge, Rh, and La, prepared from mono-element standard solutions, was used as an internal standard to monitor and correct any instrumental drift.

All calibration curves showed high linearity within the calibrated range (R^2^ > 0.9994). Instrument accuracy was evaluated by measuring a certified reference material (TMDA 64.3, fortified water), which exhibited a range of 90-105%. The limit of determination (LOD) and quantification (LOQ) for each element was calculated as the mean of the method blank concentration plus three and ten standard deviations, respectively. Some elements (Ti, V, Co, Ni, Se, Mo, Ag, Sn, Tl) exhibited concentrations lower than the LOQ, and thus were excluded from subsequent data analysis. The accuracy of the method was estimated through spike-and-recovery experiments, which yielded a range of 89-108%. This result confirms that no significant element loss occurred during the digestion procedure. The precision, determined as the repeatability of independent replicates of the same sample prepared and measured on different days, was found to be, on average, less than 10%. In conclusion, these results demonstrated that the employed method yielded satisfactory data for the objectives of this study.

### 2.3 Sr isotope analysis

To eliminate elements that could potentially cause isobaric interferences during the Sr isotope analysis (specifically Ca and Rb), digested samples underwent a Sr/matrix separation through a Sr-specific resin (SR-B100-S, particle size 50-100 µm, Triskem International, Bruz, France), in accordance to the methods proposed by Swoboda et al. (2008) and Durante et al. (2013), with minor modifications. The resin suspension was prepared by dispersing 10 g of resin in 50 mL of HNO_3_ (1% w/w), agitated for 30 min using an overhead shaker, and allowed to rest overnight. After discharging the supernatant, the resin was refilled with fresh HNO_3_ (1% w/w) to 100 mL. Before each use, the resin suspension was agitated for 30 min. For the Sr-matrix separation, 2 mL of resin was transferred onto SPE columns and washed with 2 mL of distilled water. The separation procedure included the following steps: i) resin activation with 6 mL of 8 M HNO_3_, ii) sample loading (8 M HNO_3_), iii) washing with 6 mL of 8 M HNO_3_, and iv) Sr recovery with 5 mL of distilled water. The efficiency of the Sr/matrix separation was verified by ICP-MS, demonstrating Sr recoveries higher than 85 % and negligible Ca and Rb concentrations.

The ^87^Sr/^86^Sr ratio was measured with a double-focusing multicollector inductively coupled plasma mass spectrometer (MC ICP-MS, Neptune Plus™, Thermo Scientific, Bremen, Germany) with a forward Nier-Johnson geometry. The multicollector, equipped with nine Faraday cups (a fixed central cup and eight movable ones), enabled the simultaneous measurement of the following ion beams: ^82^Kr, ^83^Kr, ^84^Sr, ^85^Rb, ^86^Sr, ^87^Sr and ^88^Sr, with ^86^Sr measured on the central cup.

To measure the ^87^Sr/^86^Sr ratio, wet plasma conditions were applied for samples with high Sr levels (Sr concentration adjusted to 200 ng/mL), while dry plasma conditions were applied to samples with low Sr levels (Sr concentration adjusted to 20 ng/mL). The data were acquired in static mode and at low resolution. The NIST SRM 987 certified reference material was measured at regular intervals in each sequence, bracketed with blank solutions. The ^87^Sr/^86^Sr ratio measured on the NIST SRM 987 within the experimental period was found to agree with both the certified and the “generally accepted value” (Stein et al., 1997), with an average precision of 0.0014% (RSD). The raw data were processed through blank subtraction, mathematical correction for the isobaric interference of Kr and Rb on Sr isotopes, and normalization of the ^88^Sr/^86^Sr value to the IUPAC value of 8.3752 to correct for mass bias induced by the instrument. Moreover, the results obtained on different days were all standardised to the NIST SRM 987 “generally accepted” value, with the application of a correction factor (Durante et al., 2013).

### 2.4 Statistical analysis

A first data evaluation was performed using principal component analysis (PCA). Before running the PCA model, the dataset was scaled and centred. Subsequently, a random forest (RF) was employed as a supervised learning method, whereby information regarding preexisting classification is provided. The model was validated by dividing the dataset into a training set (80% of the sample) and a test set (20% of the samples). In the context of classification, the results were presented in a confusion matrix, and the performance of the models was assessed based on their sensitivity, specificity, precision, negative predicted value, balanced accuracy, and Cohen’s kappa (McHugh, 2012; Tharwat, 2020).

The most significant variables identified in the RF models were subjected to further analysis, such as a Student’s t-test or the analysis of variance when comparisons were made between more than two groups, considering the species or the origin as grouping factors. In instances where the conditions of normal distribution and homoscedasticity were not met, the equivalent non-parametric tests (Wilcoxon or Kruskal-Wallis) were applied. To run a non-parametric factorial ANOVA, the dataset was first converted using the aligned rank transformation. The level of significance was set at p-value = 0.05, and Tukey’s HSD post-hoc test was employed for multiple comparisons. The statistical analysis was performed using the R computing environment (R Core Team, 2022, version 4.2.2).

## 3. Results

### 3.1 Exploratory data analysis

Analysing the dataset through the principal component analysis (PCA), a distinct clustering of the sample by species was evident (Fig. 2). This suggests that the primary source of variation in the dataset is associated with the different tree species. Considering the first two principal components which account ca. 40% of the total variance, the separation between tree species occurs along the first PC separating PIAB from LADE and PICE, which are then mostly separated along the second PC. The variables with the highest weight are the Ca/Mg concentration ratio, Ca and Sr concentration on PC1, and the concentration of Mg, K, and Ca/Sr concentration ratio on PC2, respectively, in order of contribution. A more detailed examination of the variables influencing the separation of tree species reveals that, on average, LADE samples exhibit low concentrations of nearly all the elements under consideration. In contrast, PIAB samples exhibited high concentrations of B, Ba, Ca, Sr, and a high Ca/Mg concentration ratio, which are comparatively low in the other two species. PICE samples, on the other hand, displayed high concentrations of Mg, Cr, Rb, along with high Ca/Sr and Rb/Sr concentration ratios.

Regarding the provenance of the trees, the distinction according to the bedrock type is more intricate, given that the discrepancies among the groups are less discernible (Fig. 2). Nevertheless, a partial separation among samples within each species group can be recognized along the first PC, although not particularly well-defined. In general, it can be observed that samples collected from forest stands grown on a gneiss, phyllite, or porphyry substrate are positively correlated with a high Rb/Sr concentration ratio and ^87^Sr/^86^Sr ratio, while samples collected from areas characterised by the presence of a dolomite or calc-schists substrate are positively correlated with high Ca and Sr concentrations and Ca/Mg concentration ratio.

**Figure 2.**
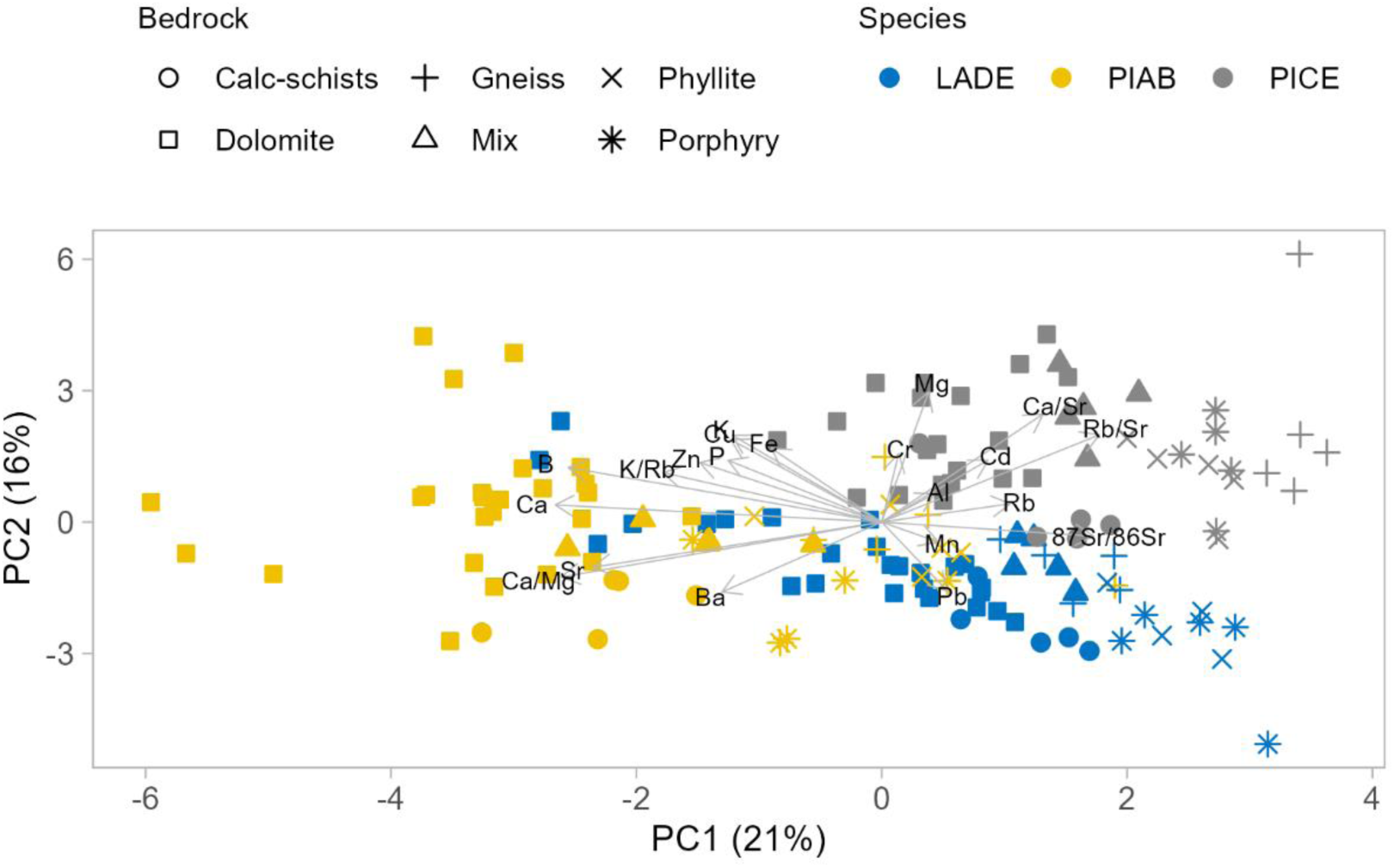
Principal component analysis (PCA) biplot of wood samples and explanatory variables (n = 20). The biplot shows the PCA scores of the explanatory variables as vectors (grey arrows) and samples (symbols). The magnitude of the vectors shows their relative contribution to each principal component (PC). The direction of the vectors indicates whether the variables are positively (in the same direction) or negatively (in opposite directions) correlated.

### 3.2 Classification model based on tree species

Based on the output of the PCA, an RF model was built by integrating multiple decision trees, enabling comprehensive classification of the samples according to tree species. The model demonstrated optimal performance with sensitivity, precision, specificity, and balanced accuracy at 100%, and a Cohen’s kappa of 1 (Table S1, Supplementary Material A).

The variable importance scores of the RF model indicated which variables from the multichemical profile were most effective in distinguishing the tree species. The mean decrease accuracy (MDA) and the mean decrease of the Gini index (MDG) were employed to ascertain the variables with the highest importance in the model development. The results indicated that the Ca concentration was the most important variable, followed by the Zn and Ba concentration (Fig. 3). For a comprehensive overview of the variable importance score for each species, please refer to Fig. S1 (Supplementary Material A). The RF partial dependence plots (Fig. S2, Supplementary Material A) indicate that Ca concentrations can be used to achieve an optimal separation between PIAB samples and PICE and LADE samples, which can be further separated by comparing their Ba concentrations, while the Zn partial dependence plot may be of limited informative value.

**Figure 3.**
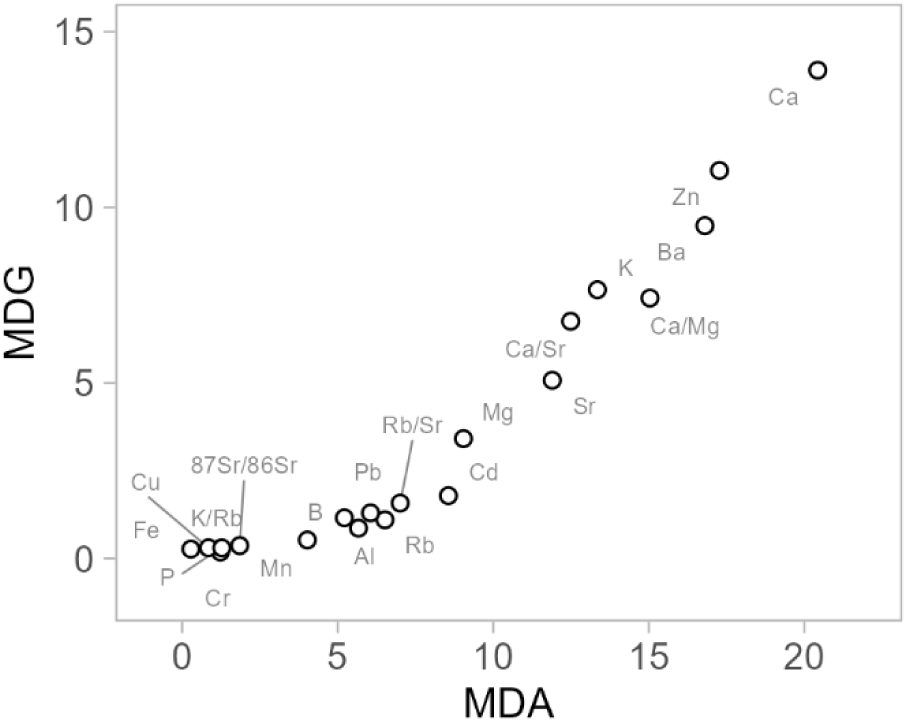
Variable importance using the mean decrease of accuracy (MDA) and mean decrease of the Gini index (MDG) measures of random forest analyses for the development of a sample classification model based on tree species.

The statistical analysis (Kruskal Wallis test) revealed that these variables can already provide a complete separation among the sample species, with significant differences among the groups detected in all three cases (Fig. 4). As indicated by the PCA outputs, the PIAB samples exhibited the highest concentrations of Ca (792 ± 170 μg g^-1^), Zn (9.7 ± 2.4 μg g^-1^) and Ba (10.4 ± 6.7 μg g^-1^). PICE samples exhibited intermediate values for both Ca and Zn (382 ± 73 μg g^-1^and 6.8 ± 1.7 μg g^-1^, respectively), while Ba concentrations was very low (<0.5 μg g^-1^, on average). The LADE samples exhibited low concentrations of Ca and Zn (262 ± 96 μg g^-1^and 3.2 ± 6.8 μg g^-1^, respectively) and intermediate Ba concentrations (5.3 ± 4.3 μg g^-1^). The clear distinction in element concentrations allowed the combination of these three variables to form the basis of a robust sample classification.

**Figure 4.**
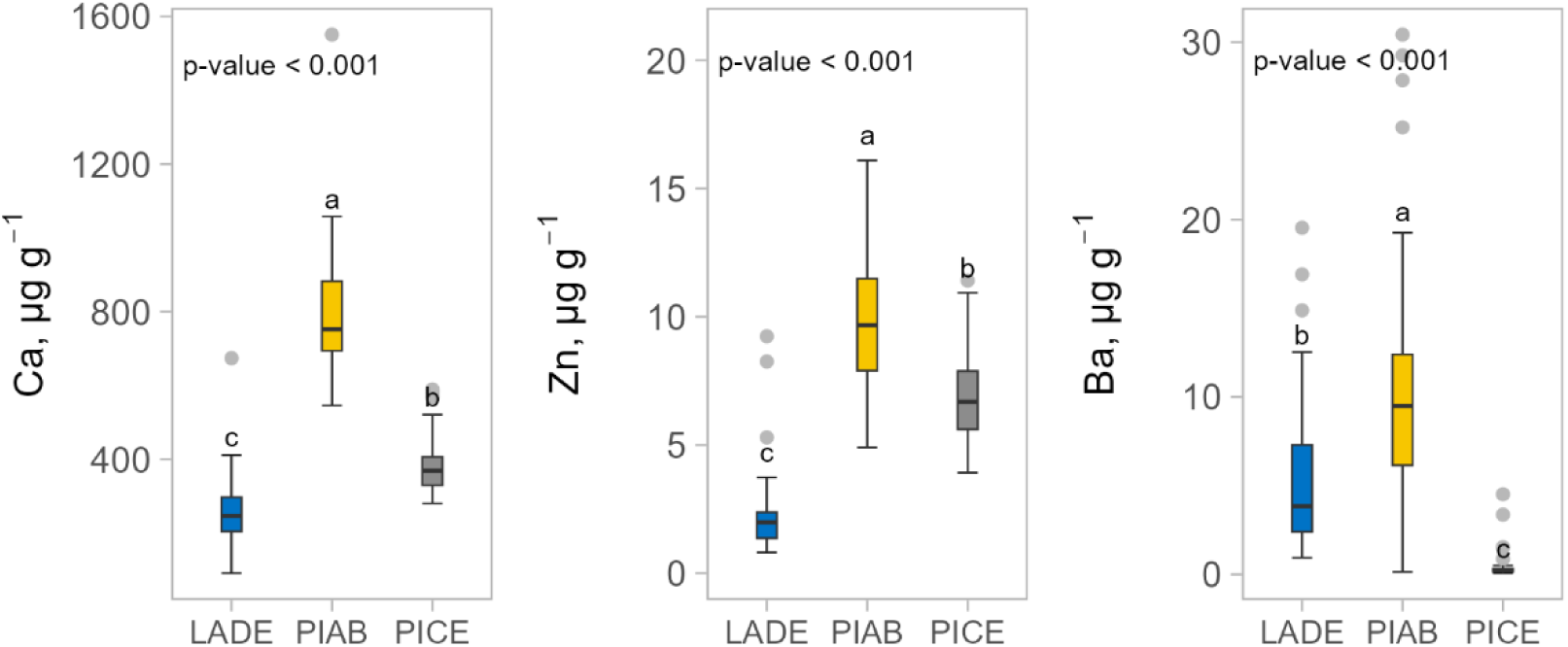
Mean concentrations of Ca, Zn, and Ba measured in the wood samples and grouped by tree species.

### 3.3 Classification models based on tree geographical origin

#### 3.3.1 Comparison among sampling sites characterized by different bedrock types

Regardless of the tree species, it is noteworthy to understand which variables are mostly bedrock-related and can be used to identify the different forest stands. A second RF model was then developed, starting from the entire dataset and setting the bedrock type as the grouping factor. The output of the model indicates that the variables with the highest importance score combining the MDA and MDG are the ^87^Sr/^86^Sr ratio, the B concentration, and the K/Rb concentration ratio (Fig. 5, see Fig. S3 in Supplementary Material A for the variable importance score plot divided by bedrock type). The performance estimators of the RF model indicate that a 100% sensitivity, specificity, precision, balanced accuracy, and Cohen’s kappa of 1 was reached during the training phase. In the validation phase, slightly lower but still satisfactory scores were obtained, due to a misclassification of a sample belonging to the calc-schist group and assigned to the dolomite group (96% sensitivity, 99% precision and specificity, 97% balanced accuracy, and 0.95 Cohen’s kappa) (Table S2, Supplementary Material A). An example of a decision tree extrapolated from the RF model is provided in Fig. S4 (Supplementary Material A).

**Figure 5.**
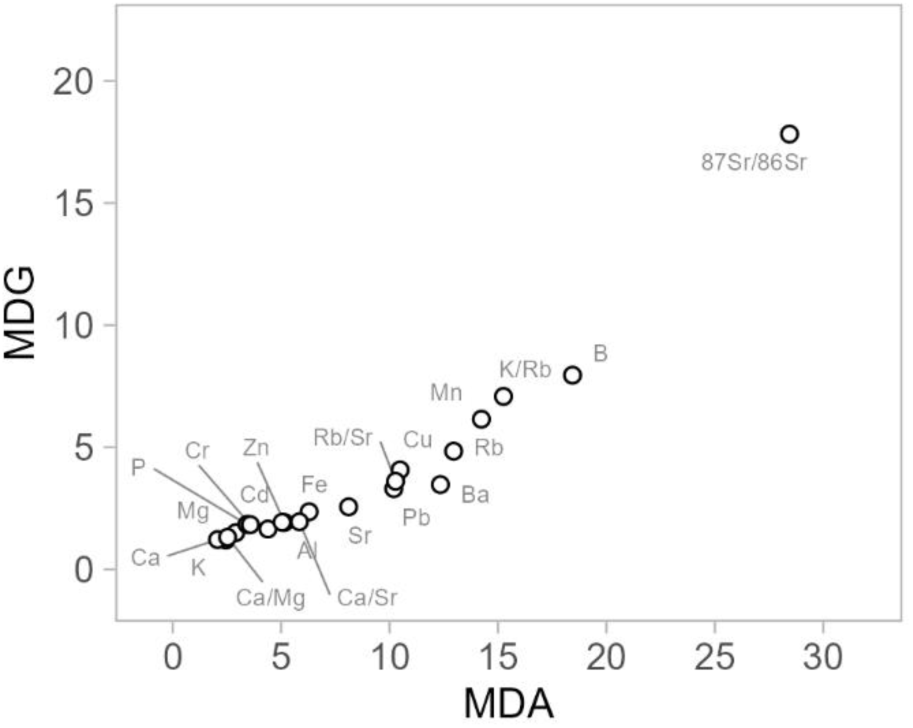
Variable importance using the mean decrease of accuracy (MDA) and mean decrease of the Gini index (MDG) measures of random forest analyses for the development of a sample classification model based on tree origin, using the bedrock type as grouping factor.

As indicated by the PCA, the sample multielement profile is found to be significantly influenced by the species. It is therefore important to understand how the relative importance of species-specific variables varies. To this end, the dataset was divided according to the tree species, and a RF model was constructed for each of them (LADE, PIAB, and PICE). A review of the variables with the highest MDA and MDG, revealed a consistent pattern across species (Fig. 6). The top three variables in terms of importance score were found to be the ^87^Sr/^86^Sr ratio and the B concentration, together with the K/Rb concentration ratio for LADE samples and Mn concentration for PIAB and PICE samples. These findings are in agreement with those obtained without the subdivision per species (Fig. 5). This suggests that the isotope ratio and concentration of these elements were more closely associated with the bedrock layer characteristic of each forest site and less dependent on the tree species. Indeed, in the RF model based on the species type, they all demonstrated a relatively low level of importance (Fig. 3). In general, the single RF models permitted a clear separation of the samples. All the performance indicators reached 100% in the training phase for LADE, PIAB, and PICE. The classification ability was slightly lower in the validation phase for PIAB and PICE, with the misclassification of a few samples affecting the performance (Tables S3, S4, and S5, Supplementary Material A). Misclassification occurred between a sample assigned to the dolomite group instead of the calc-schists group, to the gneiss group instead of the porphyry group, and to the porphyry group instead of the phyllite group.

**Figure 6.**
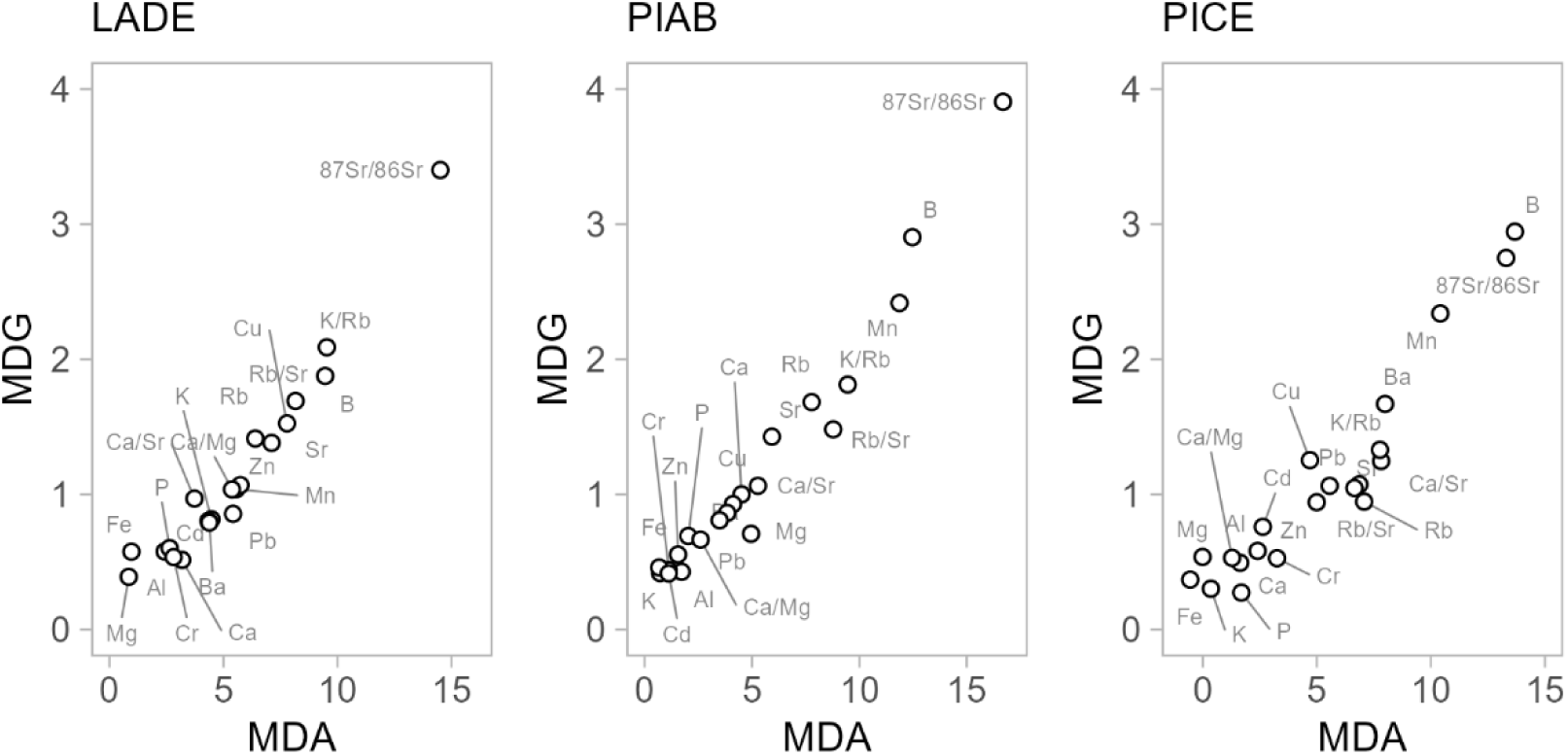
Variable importance using the mean decrease of accuracy (MDA) and mean decrease of the Gini index (MDG) measures of random forest analyses for the development of a sample classification model based on tree origin and keeping separated the three tree species.

#### 3.3.2 Comparison among sampling sites characterized by the same bedrock type

So far, the comparative analysis has focused on forest stands growing on different bedrock substrates. Nevertheless, forest sites that are geographically distant from one another may exhibit a similar bedrock type. Therefore, we sought to ascertain whether our multichemical methodology could yield sufficient data to facilitate the classification of the samples according to their geographical provenance in this case too. Regardless of the species, results demonstrated that it is still possible to identify characteristic patterns among variables that depend on the provenance. In particular, the ^87^Sr/^86^Sr ratio made a significant contribution to the classification of the samples, ranking first in terms of importance when combining the MDA and MDG scores. This was followed by the K/Rb concentration ratio, and Rb (Fig. 7 and S5 in Supplementary Material A for the variable importance score plot divided per sampling site). The model demonstrated optimal performance, exhibiting a sensitivity, specificity, precision, and balanced accuracy of 100% and a Cohen’s kappa of 1 during both the training and validation phases (Table S6, Supplementary Material A). An example of a decision tree extrapolated from the RF model is reported in Figure S6 (Supplementary Material A). Since samples from three distinct species were collected at each sampling site, the RF model was applied to the data specific to each tree species. However, this resulted in a significant reduction in the number of samples included in the model due to the introduction of an important degree of dataset fractionation. The comparison of the MDA and MDG importance scores confirmed the variables with the greatest impact, with the ^87^Sr/^86^Sr ratio among those variables identified as having the greatest influence, together with certain element concentrations and concentration ratios. Notwithstanding the limited number of samples, the models exhibited commendable performance, except for the LADE model in the validation phase, which was affected by a considerable number of sample misclassifications. Further details can be found in Fig. S7 and Tables S7, S8, and S9 in Supplementary Material A.

**Figure 7.**
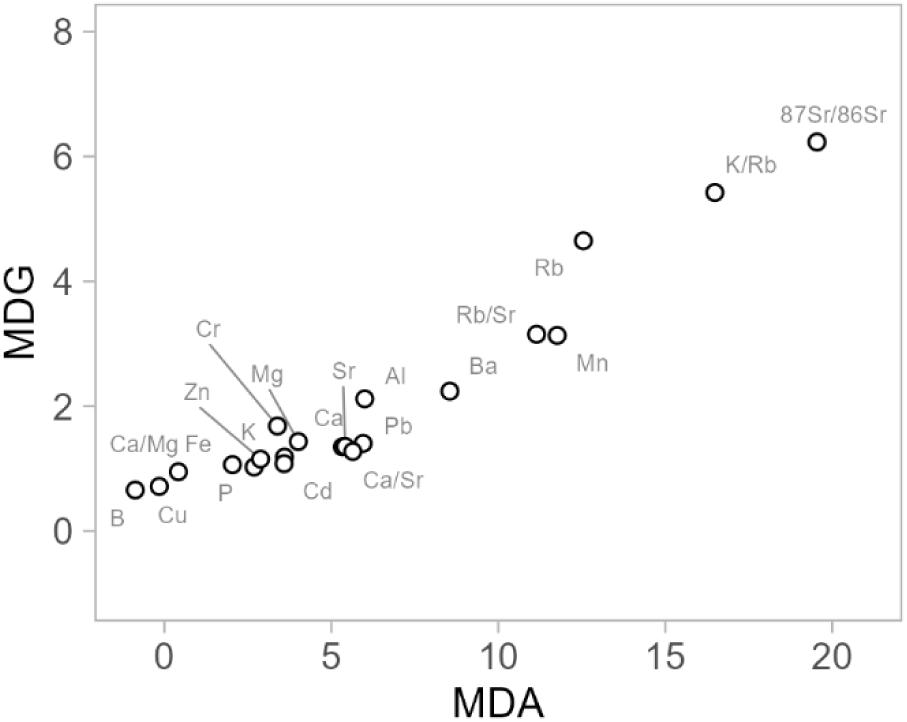
Variable importance using the mean decrease of accuracy (MDA) and mean decrease of the Gini index (MDG) measures of random forest analyses for the development of a sample classification model based on tree origin and comparing different sampling sites characterized by the same bedrock types.

As indicated by the RF models, the ^87^Sr/^86^Sr ratio has been identified as one of the most crucial variables for sample classification based on their provenance, but not for species recognition. A two-factor statistical analysis, with species and bedrock as the grouping factors, revealed that no significant difference could be attributed to either the species or the interaction between species and bedrock type, with the ^87^Sr/^86^Sr ratio demonstrating high interspecies homogeneity within individual forest stands (Fig. 8). Conversely, the observed variability is primarily attributable to the bedrock type. A further investigation of the stands belonging to the same bedrock type (dolomite) indicates that despite a very low variability of the ^87^Sr/^86^Sr ratio, with all the samples falling in a narrow range of values (0.7078-0.7097), statistically significant differences among sampling sites are still present and recognizable (Fig. 8).

**Figure 8.**
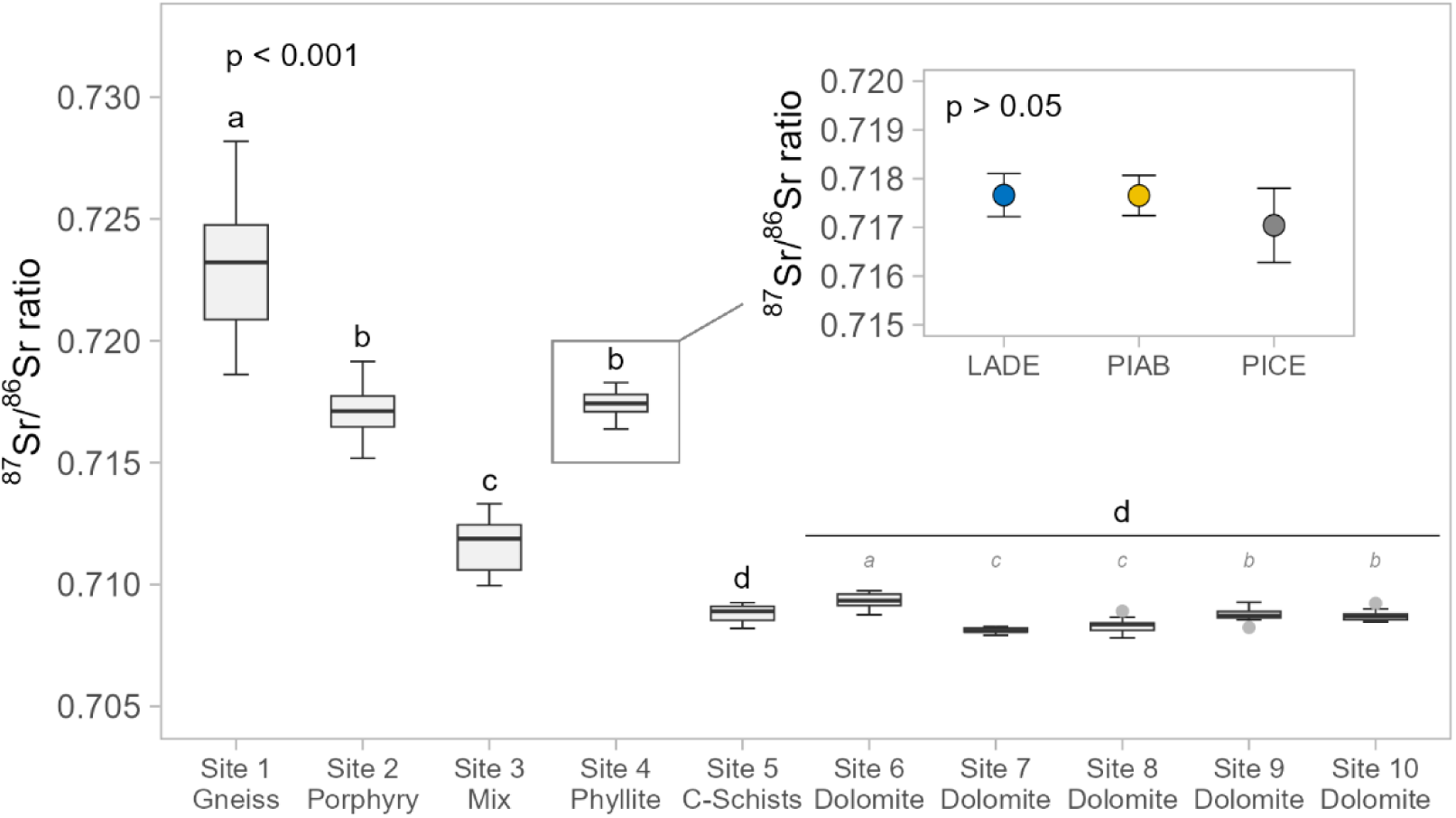
Variability of the ^87^Sr/^86^Sr ratio grouping samples according to the sampling sites. Details about the bedrock type for each sampling site are also included. Different letters denote significant differences among sampling sites. Letters in italics refer to the comparison among sampling sites of the dolomite group. In the zoomed inset the comparison among the average ^87^Sr/^86^Sr ratios measured in the samples of the different tree species collected within a single sampling site is provided.

## 4. Discussion

### 4.1 The multielement profile for timber provenancing and authenticity verification

The multielement analysis is a powerful technique that allows the rapid and simultaneous determination of several analytes within a single sample, and it is widely applied for traceability studies (Boeschoten et al., 2023b; Deklerck, 2023; Deklerck et al., 2020). The multielement composition of wood is predominantly influenced by soil chemistry, although species-specific factors also play a role (Amais et al., 2021; Cadahía et al., 1998; Van Sundert et al., 2018). For this, results of the multielement analysis can be used to group samples according to their provenance since they exhibit a distinctive regional fingerprint and, at the same time, the chemical profile of different species growing on the same substrate could be discernible (Asare and Hejcman, 2022; Boeschoten et al., 2023b). In a practical application, the macroscopic identification of the three tree species under consideration may be achieved through a non-destructive inspection, as these species exhibit distinctive anatomical wood characteristics. Nonetheless, this identification process is often critical and related to human expertise (Barmpoutis et al., 2018), hence the availability of novel approaches based on wood intrinsic characteristics is desirable.

The results of our study confirm that the classification of wood according to tree species or tree origin is possible when different groups of elements are considered. The species classification model identified mineral nutrients, including Ca, Zn, and K, as the most significant elements in wood. Additionally, it highlighted the importance of non-essential elements, such as Ba and Sr (Figure 3). While we can expect that the three tree species under examination will exhibit differing nutrient requirements and, consequently, selective absorption of nutrients, the observed difference in Ba and Sr absorption may be attributed to their chemical properties. Indeed, as both these elements belong to the same chemical group as Ca, we hypothesise that their absorption may be, to a certain extent, related to that of the major nutrient and therefore species-specific.

Irrespective of the species under consideration, other element concentrations exert a significant influence on the development of origin classification models (Fig. 5 and 7). These include micronutrients such as B and Mn, as well as non-essential elements such as Rb (including the K/Rb and Rb/Sr ratio). It seems reasonable to posit that the soil chemical features and mineral composition determine the overall bioavailability of these elements, whose uptake is less controlled by species-dependent mechanisms. It is noteworthy to highlight that the element ratios of K/Rb and Rb/Sr measured in the tree samples act as origin markers. Their variability from site to site derives from the specific composition of local minerals, hence providing information about the growing area. In particular, Rb is primarily present in K-bearing minerals (e.g. micas and potassium feldspars), as it can fit the K^+^ sites, and in igneous rocks its concentration is higher in the minerals formed in the late stages of fractional crystallization (Faure and Powell, 1972). On the contrary, the Sr distribution in minerals depends on its ability to replace Ca ions in Ca-bearing minerals or K ions in K-bearing minerals, and its concentration generally decreases with increasing magma fractionation. In agreement with these statements, we found on average the highest K/Rb ratio and the lowest Rb/Sr ratio in tree samples from the dolomitic sites, while the opposite trend characterized samples from sites with the presence of gneiss, porphyry, and phyllite in the rock substrate. To gain a more comprehensive overview a deeper geo-lithological investigation of each site would be necessary.

Finally, it should be mentioned that wood is a highly complex matrix. The multielement composition of wood may vary depending on the tree tissue sampled, and a temporal trend may also be observed over the lifetime (Boeschoten et al., 2023b). In the present study, a wood core was collected at breast height comprising the tree rings and the pith. Although there is evidence that multielement composition may exhibit temporal dynamics due to changes in the cation bioavailability in the soil, which are reflected in the multielement profile of different tree rings (Rice et al., 2022), there is a paucity of information regarding the variability along the tree height. This aspect was not evaluated in the present study, but it may warrant further investigation, particularly considering the potential applicability of this approach for precision provenancing purposes.

### 4.2 A crucial origin tracer: the Sr isotope ratio

The ^87^Sr/^86^Sr ratio played a major role in differentiating samples according to their provenance and, in all the models developed in the present study, it was among the top variables in terms of importance score, both when comparing samples from forest sites characterised by different bedrocks and within the same bedrock type.

The mean value and variability observed for the ^87^Sr/^86^Sr ratio within each stand are linked to the local geo-lithological characteristics of the area, irrespective of their geographical location. Trees absorb Sr predominantly through their root system, drawing it from the bioavailable Sr pool present in the soil. The forest sites included in this study are natural areas for which it is reasonable to conclude that this Sr fraction was gradually released from the minerals present in the bedrock through mechanisms of differential weathering. The ^87^Sr/^86^Sr ratio that is currently measured is a weighted average of that of the bedrock minerals, depending on their Rb/Sr ratio and age (Drouet et al., 2007). While the absorption of Sr by trees can be affected by several factors, including environmental conditions, soil physical and chemical properties, and species-dependent mechanisms, its isotope composition remains consistent during root uptake and distribution to the tree, as fractionation processes can be neglected when considering the precision level of this study (Marchetti et al., 2017). For this, the ^87^Sr/^86^Sr ratio is particularly effective in studies focused on sample provenancing and authenticity. Given the distinctive and diverse pedological, geological and lithological characteristics of the region under study (Gruber et al., 2019), it is reasonable to expect considerable variability in the ^87^Sr/^86^Sr ratio between forest sites and a general alignment between the findings of this study and those reported in the existing literature for these bedrock types.

Site n. 1 is located in the Val Venosta (South Tyrol), a mountainous area characterised by basement rocks with a Variscan metamorphic imprint of medium to high grade. The predominant minerals in this area are gneiss, micaschists, and quartz (Martin et al., 2009). The ^87^Sr/^86^Sr ratio, as determined from wood samples and deriving from these minerals, exhibits relatively high isotope ratios and considerable variability (0.7232 ± 0.0030). This is presumed to be associated with soil development on heterogeneous coarse-grained sediments. Similar findings have been documented in French regions with a high concentration of orthogneiss rocks, with reported ratios ranging from 0.715 to 0.721 (Willmes et al., 2018).

Site n. 2 is situated at the south-east border between South Tyrol and Trentino. The ^87^Sr/^86^Sr ratio determined in wood samples exhibits a considerable range of values, spanning from 0.7152 to 0.7192. This finding is consistent with the geo-lithological characteristics of the region. In this area, Permian vulcanites derived from the Atesino Volcanic Group are the dominant geological feature. These rocks are mainly porphyry with a high silicate content.

As illustrated on the geological maps, site n. 3 is situated at the boundary between disparate geological units, namely Permian vulcanites, Permo-Jurassic sedimentary succession, and glacial deposits of the Lower Triassic. The ^87^Sr/^86^Sr ratio measured on wood samples varies between 0.7100 and 0.7133, slightly higher compared to the typical values observed in seawater derived minerals, for which less enriched values are usually reported (Wierzbowski, 2013). This finding corroborates the hypothesis that the soil derives from a more complex geological basement.

A monotonous low-grade variscan metamorphic basement, namely the Bressanone quartz phyllite, characterises site n. 4. The ^87^Sr/^86^Sr ratio of wood samples from the site ranged from 0.7164 to 0.7183. These values are comparable to those reported for multiple rock and mineral samples collected in the Central Pyrenees, in an area characterized by graphitic phyllite (from 0.7147 to 0.7171) (McCaig et al., 2000).

The bedrock underlying site n. 5 is composed mainly of calc-schists, namely argillaceous limestone containing calcite as a substantial component and with a schistose structure, with ophiolites. The ^87^Sr/^86^Sr ratio of wood samples falls within a narrow range of values, between 0.7082 and 0.7093, which is consistent with the ratios observed in areas rich in carbonates. This is likely due to their homogeneity and relatively higher solubility than non-carbonate bearing sources (Bacher et al., 2023; Horacek, 2022; Ladegaard-Pedersen et al., 2022). Similarly, comparable values were observed in the wood samples obtained from all the sampling sites situated in the neighbouring regions of South Tyrol (sites 6 to 10, ranging from 0.7078 to 0.7097). The sites are distinguished by a carbonate/dolomite-type bedrock, which belongs to a diverse range of sedimentary successions that occurred between the Paleozoic and Mesozoic eras. These formations are generally homogeneous and exhibit low ^87^Sr/^86^Sr ratios (Willmes et al., 2018).

It is also noteworthy to highlight that no significant difference was found among wood samples from different species collected at the same sampling site. This is consistent with the results reported by Erban Kochergina et al. (2021), who conducted a comparative analysis of oak and pine samples collected from the same sampling sites. From a provenance and authenticity perspective, this allows the creation of regional databases for the Sr isotope ratio that can be used as a reference regardless of the species under consideration. This represents a significant advantage of this marker in comparison to other geographical indicators that are typically employed in traceability studies, including the multielement profile and the stable isotope ratio of light elements. In the latter cases, indeed, a significant species effect has been observed, necessitating the creation of species-specific reference datasets (Amais et al., 2021; Boeschoten et al., 2023a). Furthermore, both the stable isotope ratio of light elements and the multielement profile may be subject to year-to-year fluctuations (Boeschoten et al., 2023a, 2023b; Gori et al., 2018; Rice et al., 2022), thereby complicating the process of proper sampling and material preparation. Conversely, the ^87^Sr/^86^Sr ratio exhibits minimal variability from outer growing rings to the pith of the tree trunk, falling within the intra-site variability range (see Supplementary Material B).

### 4.3 The strength of a multichemical approach and predictive models

It can be observed that none of the considered variables under consideration can provide sufficient information to enable the classification of an unknown sample, particularly when the geographical origin is taken into account. The principal benefit of a multichemical methodology is that it permits the examination of a greater number of variables, thereby facilitating the utilisation of chemometrics tools. The application of supervised methods allows the evaluation of the influence of the different factors on the variability of the dependent variable, thereby enabling the extrapolation of general principles that can be employed to make predictions, resulting in a markedly more accurate classification output (González-Domínguez et al., 2022).

This study integrates the findings of the multielement analysis with those of the Sr isotope ratio analysis, demonstrating that this multichemical approach can provide sufficient information to develop classification models that consider both the sample species and the sample origin as grouping factors. While several studies have demonstrated the feasibility of this approach for food provenancing (Aguzzoni et al., 2021; Jandric et al., 2021; Monti et al., 2022; Zuliani et al., 2020), less information was available regarding forestry products. The RF algorithm demonstrated that classification models with varying spatial resolutions can be generated, shifting the focus from a broader perspective (different bedrock types) to a highly detailed stand differentiation (within the same bedrock type). This approach consistently yielded excellent performance, with balanced accuracy exceeding 90% in most cases.

The present study encompasses a restricted yet diverse mountain area in the Alpine region, characterised by a varied bedrock composition. The data obtained from this study provide sufficient information to support the use of this approach to certify different timber production areas. This could be of significant importance in supporting local certified productions in mountain communities, for which the timber economy is of vital importance (Gori et al., 2018). Providing objective evidence of the wood provenance may facilitate the revitalisation of the compartment, the promotion of sustainable forest management, and the guarantee of the product’s origin. The geographic origin of timber is a crucial aspect of international trade and it is essential to ensure that it is conducted in compliance with the current legislation (Lowe et al., 2016). This is pivotal considering that the demand for wood products is accompanied by a dramatic incidence of illegal logging, which is diffused in both tropical and non-tropical regions, causing severe deforestation and a loss of biodiversity in several regions worldwide (Dormontt et al., 2015; Lowe et al., 2016). The current approach primarily focuses on the development of international regulations, which are often complemented by techniques such as paper-based certificates and barcoding. However, these methods are inadequate for verifying compliance with declarations, particularly with regard to species identification and geographic origin (Dormontt et al., 2015; Lowe et al., 2016). In this context, the adoption of scientific methods based on intrinsic wood properties, such as a multichemical approach as we presented here, can enhance forensic timber identification and bolster the integrity of certification schemes (Boeschoten et al., 2023b; Gori et al., 2018). Furthermore, the development of sample classification tools to authenticate unknown samples can support the activity of control bodies, particularly if international databases pertaining to the legal supply chain are established, as has been done for other commodities, such as wine (Schlesier et al., 2009). Ongoing efforts are needed to facilitate the transition from research to forensic tools. This process will require collaboration among stakeholders and authorities. Nevertheless, studies such as this one contribute to the advancement of knowledge regarding the potential of chemical markers in provenancing and authenticity studies, to make a step forward toward an integrated approach to address the issue of the illegal timber trade.

## CRediT authorship contribution statement

**Agnese Aguzzoni:** Conceptualization, Data curation, Formal analysis, Investigation, Methodology, Visualization, Writing – original draft, Writing – review & editing. **Francesco Giammarchi:** Conceptualization, Investigation, Formal analysis, Writing – original draft, Writing – review & editing. **Ignacio A. Mundo:** Investigation, Formal analysis, Visualization, Writing – review & editing. **Giulio Voto:** Methodology, Writing – review & editing. **Giustino Tonon:** Conceptualization, Funding acquisition, Project administration, Supervision. **Werner Tirler:** Conceptualization, Funding acquisition, Project administration, Supervision, Writing – review & editing. **Enrico Tomelleri:** Conceptualization, Supervision, Formal analysis, Writing – original draft, Writing – review & editing. All authors have read and agreed to the published version of the manuscript.

## Acknowledgments

This paper is dedicated to the memory of Professor Giustino Tonon, who passed away on 7 July 2021. A special thanks goes to the Forestry Service of the Province of Bolzano/Bozen and the neighbouring regions for their valuable support during the sampling campaign. The authors would like to express their gratitude to Raffaela Dibona and Marco Carrer of the University of Padova for facilitating the sampling at Passo Tre Croci, to Josef Walch of the Tyrol Forestry Service for accompanying us in the sampling at Hahntennjoch, to the Polizia of Livigno for authorising the sampling at Alpisella and to the Consorzio Forestale “Pizzo Camino” for allowing the sampling at Borno. Thanks to Christian Grummer, Maurizio Ventura and Lucio Dorigatti of the Free University of Bolzano for their collaboration in the sampling at Passo Tre Croci, Livigno and Borno. We would also like to thank Christian Grumer, Maurizio Ventura and Lucio Dorigatti of the Free University of Bolzano for their collaboration in the samplings conducted at Passo Tre Croci, Livigno and Borno.

