## Supplementary Material A for "Enhancing timber traceability via multielement and strontium isotope ratio: An example from the Eastern Alps"

**Table S1.** Confusion matrix for the random forest (RF) model for both the training and validation phase using the species as grouping factor. Performance metrics were calculated according to the equations reported in literature (McHugh, 2012; Tharwat, 2020).

| | **Model training True species** | | | | **Model validation True species** | | | |
|---|---|---|---|---|---|---|---|---|
| **Predicted species** | **LADE** | **PIAB** | **PICE** | **Average** | **LADE** | **PIAB** | **PICE** | **Average** |
| LADE | 41 | 0 | 0 | | 8 | 0 | 0 | |
| PIAB | 0 | 42 | 0 | | 0 | 8 | 0 | |
| PICE | 0 | 0 | 31 | | 0 | 0 | 14 | |
| Sensitivity | 100% | 100% | 100% | 100% | 100% | 100% | 100% | 100% |
| Specificity | 100% | 100% | 100% | 100% | 100% | 100% | 100% | 100% |
| Precision | 100% | 100% | 100% | 100% | 100% | 100% | 100% | 100% |
| Negative predicted value | 0% | 0% | 0% | 0% | 0% | 0% | 0% | 0% |
| Balanced accuracy | 100% | 100% | 100% | 100% | 100% | 100% | 100% | 100% |
| Cohen’s kappa | | | | 1 | | | | 1 |

LADE, European larch (*Larix decidua* Mill.); PIAB, Norway spruce (*Picea abies* Karst.); PICE, Swiss stone pine (*Pinus cembra* L.).

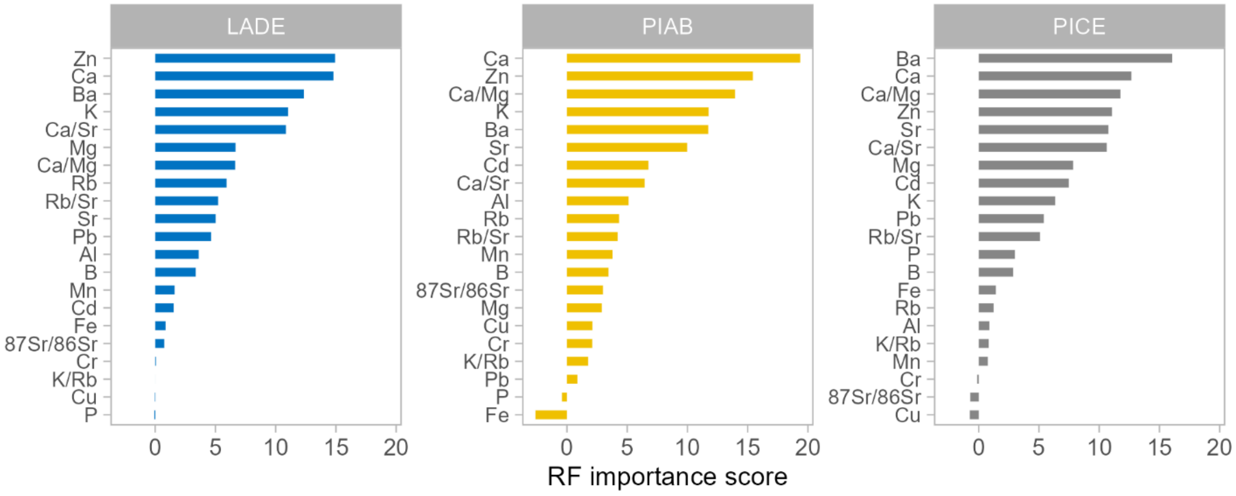


**Figure S1.** RF importance score of each variable for the model developed using the species as grouping factor.

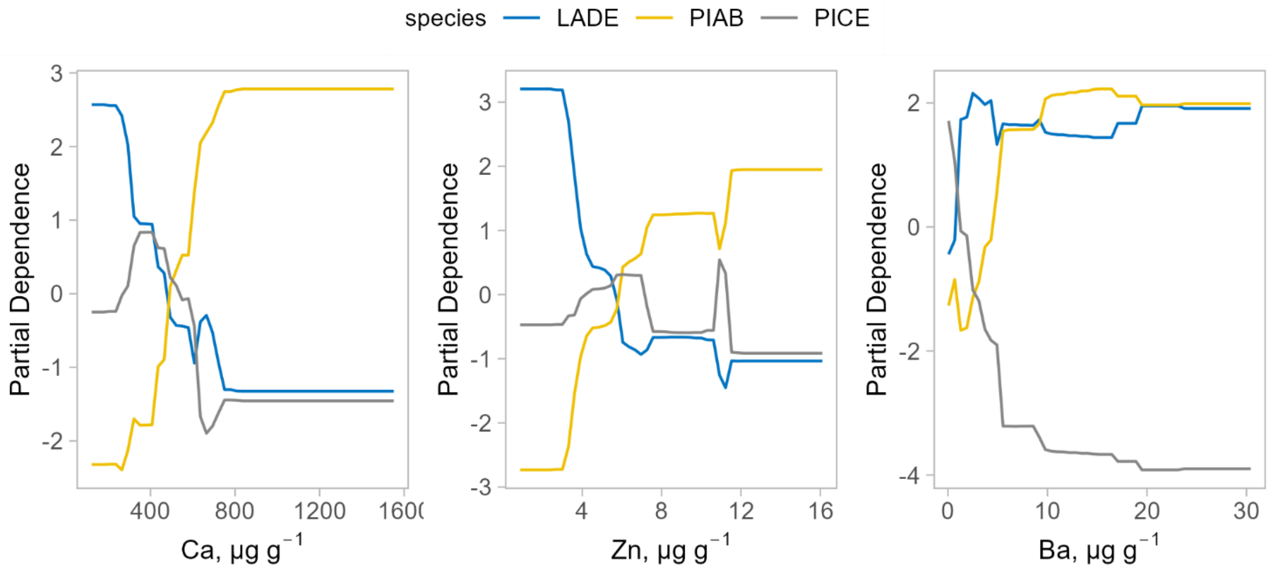


**Figure S2.** Partial dependence plots for the three elements with the highest importance score relative to the RF model developed using the species as grouping factor.

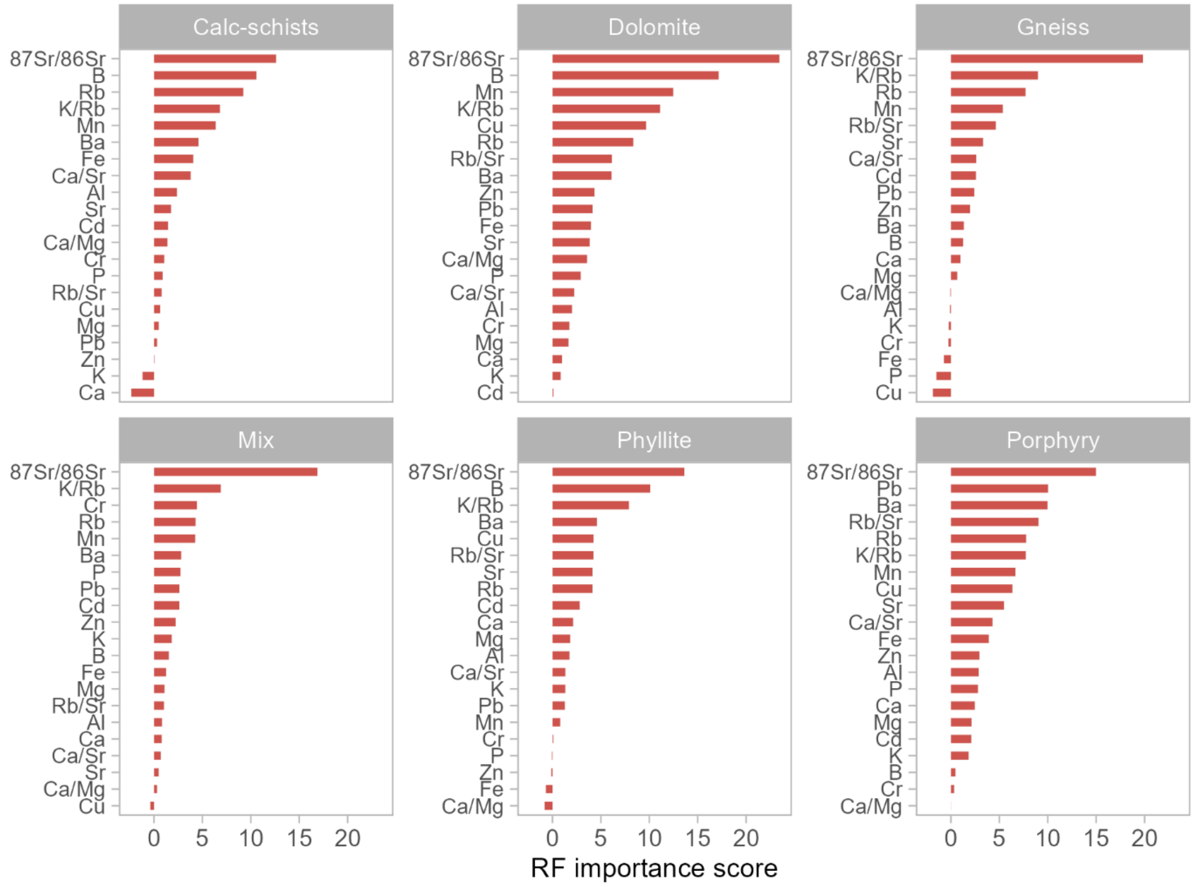


**Figure S3.** RF importance score of each variable for the model developed using the bedrock type as grouping factor.

**Table S2.** Confusion matrix for the random forest (RF) model for both the training and validation phase using the bedrock type as grouping factor. The dataset for the model development included all the samples, regardless of the species. Performance metrics were calculated according to the equations reported in literature (McHugh, 2012; Tharwat, 2020).

| | **Model training True origin** | | | | | | | **Model validation True origin** | | | | | | |
|---|---|---|---|---|---|---|---|---|---|---|---|---|---|---|
| **Predicted origin** | **Calc-schists** | **Dolomite** | **Gneiss** | **Mix** | **Phyllite** | **Porphyry** | **Average** | **Calc-schists** | **Dolomite** | **Gneiss** | **Mix** | **Phyllite** | **Porphyry** | **Average** |
| Calc-schists | 10 | 0 | 0 | 0 | 0 | 0 | | 4 | 0 | 0 | 0 | 0 | 0 | |
| Dolomite | 0 | 56 | 0 | 0 | 0 | 0 | | 1 | 14 | 0 | 0 | 0 | 0 | |
| Gneiss | 0 | 0 | 11 | 0 | 0 | 0 | | 0 | 0 | 4 | 0 | 0 | 0 | |
| Mix | 0 | 0 | 0 | 12 | 0 | 0 | | 0 | 0 | 0 | 3 | 0 | 0 | |
| Phyllite | 0 | 0 | 0 | 0 | 11 | 0 | | 0 | 0 | 0 | 0 | 3 | 0 | |
| Porphyry | 0 | 0 | 0 | 0 | 0 | 14 | | 0 | 0 | 0 | 0 | 0 | 1 | |
| Sensitivity | 100% | 100% | 100% | 100% | 100% | 100% | 100% | 80% | 100% | 100% | 100% | 100% | 100% | 96% |
| Specificity | 100% | 100% | 100% | 100% | 100% | 100% | 100% | 100% | 94% | 100% | 100% | 100% | 100% | 99% |
| Precision | 100% | 100% | 100% | 100% | 100% | 100% | 100% | 100% | 93% | 100% | 100% | 100% | 100% | 99% |
| Negative predicted value | 0% | 0% | 0% | 0% | 0% | 0% | 0% | 96% | 100% | 100% | 100% | 100% | 100% | 99% |
| Balanced accuracy | 100% | 100% | 100% | 100% | 100% | 100% | 100% | 90% | 97% | 100% | 100% | 100% | 100% | 97% |
| Cohen's kappa | | | | | | | 1 | | | | | | | 0.95 |

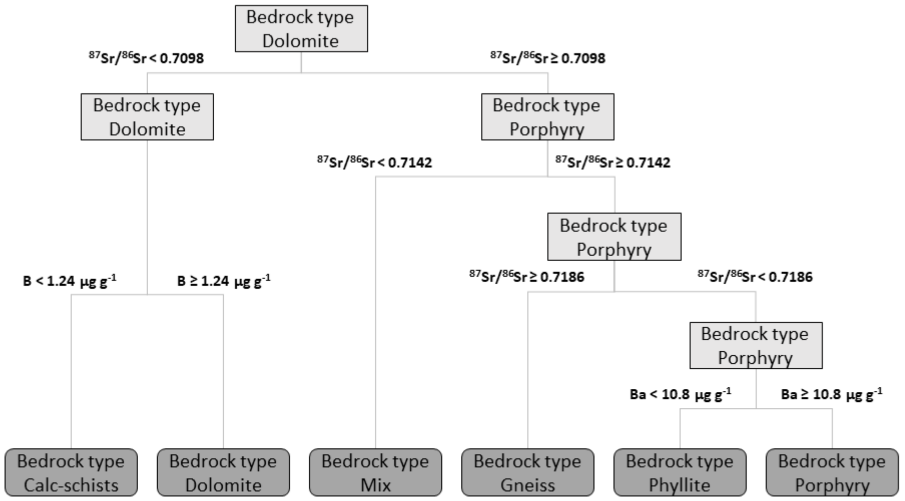


**Figure S4.** Example of a decision tree extrapolated from the random forest model for tree origin classification depending on the different bedrock types.

**Table S3.** Confusion matrix for the random forest (RF) model for both the training and validation phase using the bedrock type as grouping factor. The dataset for the model development included only samples belonging to the LADE species. Performance metrics were calculated according to the equations reported in literature (McHugh, 2012; Tharwat, 2020).

| | **Model training True origin** | | | | | | | **Model validation True origin** | | | | | | |
|---|---|---|---|---|---|---|---|---|---|---|---|---|---|---|
| **Predicted origin** | **Calc-schists** | **Dolomite** | **Gneiss** | **Mix** | **Phyllite** | **Porphyry** | **Average** | **Calc-schists** | **Dolomite** | **Gneiss** | **Mix** | **Phyllite** | **Porphyry** | **Average** |
| Calc-schists | 3 | 0 | 0 | 0 | 0 | 0 | | 2 | 0 | 0 | 0 | 0 | 0 | |
| Dolomite | 0 | 20 | 0 | 0 | 0 | 0 | | 0 | 5 | 0 | 0 | 0 | 0 | |
| Gneiss | 0 | 0 | 4 | 0 | 0 | 0 | | 0 | 0 | 1 | 0 | 0 | 0 | |
| Mix | 0 | 0 | 0 | 4 | 0 | 0 | | 0 | 0 | 0 | 1 | 0 | 0 | |
| Phyllite | 0 | 0 | 0 | 0 | 3 | 0 | | 0 | 0 | 0 | 0 | 1 | 0 | |
| Porphyry | 0 | 0 | 0 | 0 | 0 | 3 | | 0 | 0 | 0 | 0 | 0 | 2 | |
| Sensitivity | 100% | 100% | 100% | 100% | 100% | 100% | 100% | 100% | 100% | 100% | 100% | 100% | 100% | 100% |
| Specificity | 100% | 100% | 100% | 100% | 100% | 100% | 100% | 100% | 100% | 100% | 100% | 100% | 100% | 100% |
| Precision | 100% | 100% | 100% | 100% | 100% | 100% | 100% | 100% | 100% | 100% | 100% | 100% | 100% | 100% |
| Negative predicted value | 100% | 100% | 100% | 100% | 100% | 100% | 100% | 100% | 100% | 100% | 100% | 100% | 100% | 100% |
| Balanced accuracy | 100% | 100% | 100% | 100% | 100% | 100% | 100% | 100% | 100% | 100% | 100% | 100% | 100% | 100% |
| Cohen's kappa | | | | | | | 1 | | | | | | | 1 |

**Table S4.** Confusion matrix for the random forest (RF) model for both the training and validation phase using the bedrock type as grouping factor. The dataset for the model development included only samples belonging to the PIAB species. Performance metrics were calculated according to the equations reported in literature (McHugh, 2012; Tharwat, 2020).

| | **Model training True origin** | | | | | | | **Model validation True origin** | | | | | | |
|---|---|---|---|---|---|---|---|---|---|---|---|---|---|---|
| **Predicted origin** | **Calc-schists** | **Dolomite** | **Gneiss** | **Mix** | **Phyllite** | **Porphyry** | **Average** | **Calc-schists** | **Dolomite** | **Gneiss** | **Mix** | **Phyllite** | **Porphyry** | **Average** |
| Calc-schists | 3 | 0 | 0 | 0 | 0 | 0 | | 1 | 0 | 0 | 0 | 0 | 0 | |
| Dolomite | 0 | 20 | 0 | 0 | 0 | 0 | | 1 | 5 | 0 | 0 | 0 | 0 | |
| Gneiss | 0 | 0 | 4 | 0 | 0 | 0 | | 0 | 0 | 1 | 0 | 0 | 1 | |
| Mix | 0 | 0 | 0 | 4 | 0 | 0 | | 0 | 0 | 0 | 1 | 0 | 0 | |
| Phyllite | 0 | 0 | 0 | 0 | 4 | 0 | | 0 | 0 | 0 | 0 | 1 | 0 | |
| Porphyry | 0 | 0 | 0 | 0 | 0 | 3 | | 0 | 0 | 0 | 0 | 0 | 1 | |
| Sensitivity | 100% | 100% | 100% | 100% | 100% | 100% | 100% | 50% | 100% | 100% | 100% | 100% | 50% | 90% |
| Specificity | 100% | 100% | 100% | 100% | 100% | 100% | 100% | 100% | 86% | 91% | 100% | 100% | 100% | 95% |
| Precision | 100% | 100% | 100% | 100% | 100% | 100% | 100% | 100% | 83% | 50% | 100% | 100% | 100% | 87% |
| Negative predicted value | 100% | 100% | 100% | 100% | 100% | 100% | 100% | 91% | 100% | 100% | 100% | 100% | 91% | 98% |
| Balanced accuracy | 100% | 100% | 100% | 100% | 100% | 100% | 100% | 75% | 93% | 95% | 100% | 100% | 75% | 93% |
| Cohen's kappa | | | | | | | 1 | | | | | | | 0.77 |

**Table S5.** Confusion matrix for the random forest (RF) model for both the training and validation phase using the bedrock type as grouping factor. The dataset for the model development included only samples belonging to the PICE species. Performance metrics were calculated according to the equations reported in literature (McHugh, 2012; Tharwat, 2020).

| | **Model training True origin** | | | | | | | **Model validation True origin** | | | | | | |
|---|---|---|---|---|---|---|---|---|---|---|---|---|---|---|
| **Predicted origin** | **Calc-schists** | **Dolomite** | **Gneiss** | **Mix** | **Phyllite** | **Porphyry** | **Average** | **Calc-schists** | **Dolomite** | **Gneiss** | **Mix** | **Phyllite** | **Porphyry** | **Average** |
| Calc-schists | 3 | 0 | 0 | 0 | 0 | 0 | | 2 | 0 | 0 | 0 | 0 | 0 | |
| Dolomite | 0 | 17 | 0 | 0 | 0 | 0 | | 0 | 3 | 0 | 0 | 0 | 0 | |
| Gneiss | 0 | 0 | 3 | 0 | 0 | 0 | | 0 | 0 | 2 | 0 | 0 | 0 | |
| Mix | 0 | 0 | 0 | 4 | 0 | 0 | | 0 | 0 | 0 | 1 | 0 | 0 | |
| Phyllite | 0 | 0 | 0 | 0 | 3 | 0 | | 0 | 0 | 0 | 0 | 1 | 0 | |
| Porphyry | 0 | 0 | 0 | 0 | 0 | 4 | | 0 | 0 | 0 | 0 | 1 | 1 | |
| Sensitivity | 100% | 100% | 100% | 100% | 100% | 100% | 100% | 100% | 100% | 100% | 100% | 50% | 100% | 90% |
| Specificity | 100% | 100% | 100% | 100% | 100% | 100% | 100% | 100% | 100% | 100% | 100% | 100% | 90% | 100% |
| Precision | 100% | 100% | 100% | 100% | 100% | 100% | 100% | 100% | 100% | 100% | 100% | 100% | 50% | 100% |
| Negative predicted value | 100% | 100% | 100% | 100% | 100% | 100% | 100% | 100% | 100% | 100% | 100% | 90% | 100% | 98% |
| Balanced accuracy | 100% | 100% | 100% | 100% | 100% | 100% | 100% | 100% | 100% | 100% | 100% | 75% | 95% | 95% |
| Cohen's kappa | | | | | | | 1 | | | | | | | 0.89 |

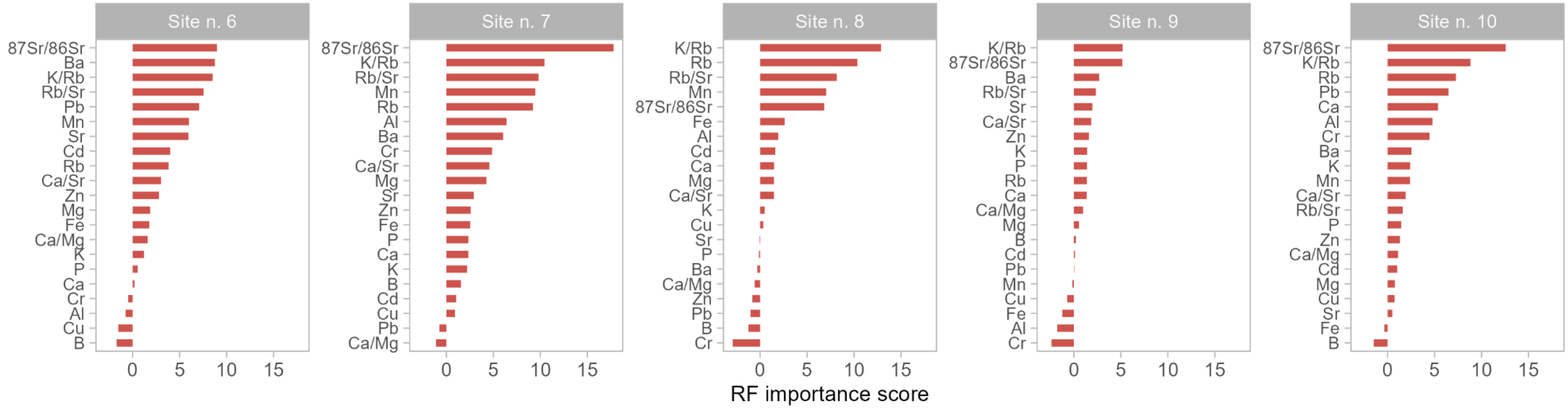


**Figure S5.** RF importance score of each variable for the model developed using the sampling site as grouping factor and considering only those characterized by dolomite bedrock. The dataset for the model development included all the samples, regardless of the species.

**Table S6.** Confusion matrix for the random forest (RF) model for both the training and validation phase comparing only samples collected in sampling sites (grouping factor) characterized by a dolomite bedrock. The dataset for the model development included all the samples, regardless of the species. Performance metrics were calculated according to the equations reported in literature (McHugh, 2012; Tharwat, 2020).

| | **Model training True origin** | | | | | | **Model validation True origin** | | | | | |
|---|---|---|---|---|---|---|---|---|---|---|---|---|
| **Predicted origin** | **Site n. 6** | **Site n. 7** | **Site n. 8** | **Site n. 9** | **Site n. 10** | **Average** | **Site n. 6** | **Site n. 7** | **Site n. 8** | **Site n. 9** | **Site n. 10** | **Average** |
| Site n. 6 | 6 | 0 | 0 | 0 | 0 | | 4 | 0 | 0 | 0 | 0 | |
| Site n. 7 | 0 | 14 | 0 | 0 | 0 | | 0 | 1 | 0 | 0 | 0 | |
| Site n. 8 | 0 | 0 | 11 | 0 | 0 | | 0 | 0 | 4 | 0 | 0 | |
| Site n. 9 | 0 | 0 | 0 | 11 | 0 | | 0 | 0 | 0 | 4 | 0 | |
| Site n. 10 | 0 | 0 | 0 | 0 | 14 | | 0 | 0 | 0 | 0 | 1 | |
| Sensitivity | 100% | 100% | 100% | 100% | 100% | 100% | 100% | 100% | 100% | 100% | 100% | 100% |
| Specificity | 100% | 100% | 100% | 100% | 100% | 100% | 100% | 100% | 100% | 100% | 100% | 100% |
| Precision | 100% | 100% | 100% | 100% | 100% | 100% | 100% | 100% | 100% | 100% | 100% | 100% |
| Negative predicted value | 100% | 100% | 100% | 100% | 100% | 100% | 100% | 100% | 100% | 100% | 100% | 100% |
| Balanced accuracy | 100% | 100% | 100% | 100% | 100% | 100% | 100% | 100% | 100% | 100% | 100% | 100% |
| Cohen’s kappa | | | | | | 1 | | | | | | 1 |

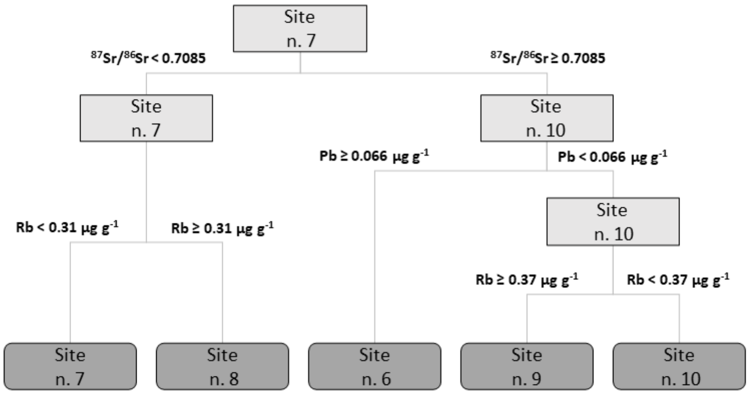


**Figure S6.** Example of a decision tree extrapolated from the random forest model for a sample classification model based on tree origin and comparing different sampling sites characterized by the same bedrock types.

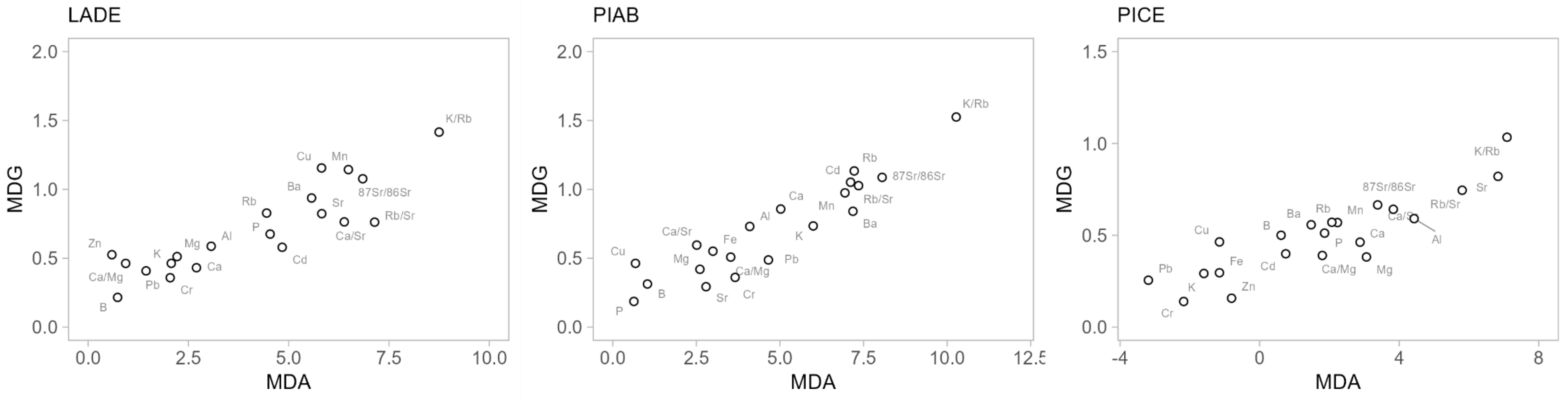


**Figure S7.** Variable importance using the mean decrease of the Gini index and mean decrease of accuracy measures of random forest analyses for the development of a sample classification model using the sampling site as grouping factor and considering only those characterized by dolomite bedrock. The dataset for the model development was subdivided according to the sample species

**Table S7.** Confusion matrix for the random forest (RF) model for both the training and validation phase comparing only samples collected in sampling sites (grouping factor) characterized by a dolomite bedrock. The dataset for the model development included only samples belonging to the LADE species. Performance metrics were calculated according to the equations reported in literature (McHugh, 2012; Tharwat, 2020).

| | **Model training True origin** | | | | | | **Model validation True origin** | | | | | |
|---|---|---|---|---|---|---|---|---|---|---|---|---|
| **Predicted origin** | **Site n. 6** | **Site n. 7** | **Site n. 8** | **Site n. 9** | **Site n. 10** | **Average** | **Site n. 6** | **Site n. 7** | **Site n. 8** | **Site n. 9** | **Site n. 10** | **Average** |
| Site n. 6 | 4 | 0 | 0 | 0 | 0 | | 1 | 0 | 0 | 0 | 0 | |
| Site n. 7 | 0 | 4 | 0 | 0 | 0 | | 0 | 1 | 0 | 0 | 0 | |
| Site n. 8 | 0 | 0 | 4 | 0 | 0 | | 0 | 0 | 1 | 2 | 0 | |
| Site n. 9 | 0 | 0 | 0 | 3 | 0 | | 0 | 0 | 0 | 0 | 1 | |
| Site n. 10 | 0 | 0 | 0 | 0 | 4 | | 0 | 0 | 0 | 0 | 0 | |
| Sensitivity | 100% | 100% | 100% | 100% | 100% | 100% | 100% | 100% | 100% | 0% | 0% | 60% |
| Specificity | 100% | 100% | 100% | 100% | 100% | 100% | 100% | 100% | 60% | 75% | 100% | 87% |
| Precision | 100% | 100% | 100% | 100% | 100% | 100% | 100% | 100% | 33% | 0% | NA | 58% |
| Negative predicted value | 100% | 100% | 100% | 100% | 100% | 100% | 100% | 100% | 100% | 60% | 83% | 89% |
| Balanced accuracy | 100% | 100% | 100% | 100% | 100% | 100% | 100% | 100% | 80% | 38% | 50% | 74% |
| Cohen’s kappa | | | | | | 1 | | | | | | 0.38 |

**Table S8.** Confusion matrix for the random forest (RF) model for both the training and validation phase comparing only samples collected in sampling sites (grouping factor) characterized by a dolomite bedrock. The dataset for the model development included only samples belonging to the PIAB species. Performance metrics were calculated according to the equations reported in literature (McHugh, 2012; Tharwat, 2020).

| | **Model training True origin** | | | | | | **Model validation True origin** | | | | | |
|---|---|---|---|---|---|---|---|---|---|---|---|---|
| **Predicted origin** | **Site n. 6** | **Site n. 7** | **Site n. 8** | **Site n. 9** | **Site n. 10** | **Average** | **Site n. 6** | **Site n. 7** | **Site n. 8** | **Site n. 9** | **Site n. 10** | **Average** |
| Site n. 6 | 4 | 0 | 0 | 0 | 0 | | 1 | 0 | 0 | 0 | 0 | |
| Site n. 7 | 0 | 4 | 0 | 0 | 0 | | 0 | 1 | 0 | 0 | 0 | |
| Site n. 8 | 0 | 0 | 4 | 0 | 0 | | 0 | 0 | 1 | 0 | 0 | |
| Site n. 9 | 0 | 0 | 0 | 3 | 0 | | 0 | 0 | 0 | 2 | 0 | |
| Site n. 10 | 0 | 0 | 0 | 0 | 4 | | 0 | 0 | 0 | 0 | 1 | |
| Sensitivity | 100% | 100% | 100% | 100% | 100% | 100% | 100% | 100% | 100% | 100% | 100% | 100% |
| Specificity | 100% | 100% | 100% | 100% | 100% | 100% | 100% | 100% | 100% | 100% | 100% | 100% |
| Precision | 100% | 100% | 100% | 100% | 100% | 100% | 100% | 100% | 100% | 100% | 100% | 100% |
| Negative predicted value | 100% | 100% | 100% | 100% | 100% | 100% | 100% | 100% | 100% | 100% | 100% | 100% |
| Balanced accuracy | 100% | 100% | 100% | 100% | 100% | 100% | 100% | 100% | 100% | 100% | 100% | 100% |
| Cohen’s kappa | | | | | | 1 | | | | | | 1 |

**Table S9.** Confusion matrix for the random forest (RF) model for both the training and validation phase comparing only samples collected in sampling sites (grouping factor) characterized by a dolomite bedrock. The dataset for the model development included only samples belonging to the PICE species. Site n. 6 was not included as it was not possible to collect PICE samples at this location. Performance metrics were calculated according to the equations reported in literature (McHugh, 2012; Tharwat, 2020).

| | **Model training<br>True origin** | | | | | **Model validation<br>True origin** | | | | |
|---|---|---|---|---|---|---|---|---|---|---|
| **Predicted origin** | **Site n. 7** | **Site n. 8** | **Site n. 9** | **Site n. 10** | **Average** | **Site n. 7** | **Site n. 8** | **Site n. 9** | **Site n. 10** | **Average** |
| Site n. 7 | 4 | 0 | 0 | 0 | | 1 | 0 | 0 | 0 | |
| Site n. 8 | 0 | 3 | 0 | 0 | | 0 | 2 | 0 | 0 | |
| Site n. 9 | 0 | 0 | 4 | 0 | | 0 | 0 | 1 | 0 | |
| Site n. 10 | 0 | 0 | 0 | 4 | | 0 | 0 | 0 | 1 | |
| Sensitivity | 100% | 100% | 100% | 100% | 100% | 100% | 100% | 100% | 100% | 100% |
| Specificity | 100% | 100% | 100% | 100% | 100% | 100% | 100% | 100% | 100% | 100% |
| Precision | 100% | 100% | 100% | 100% | 100% | 100% | 100% | 100% | 100% | 100% |
| Negative predicted value | 100% | 100% | 100% | 100% | 100% | 100% | 100% | 100% | 100% | 100% |
| Balanced accuracy | 100% | 100% | 100% | 100% | 100% | 100% | 100% | 100% | 100% | 100% |
| Cohen’s kappa | | | | | 1 | | | | | 1 |
