## Supplementary Material B for "Enhancing timber traceability via multielement and strontium isotope ratio: An example from the Eastern Alps"

### Preliminary investigation

The Sr isotope ratio is a soil-derived chemical marker largely used in traceability and authenticity studies, especially in the food sector (Baffi and Trincherini, 2016; Marchetti et al., 2017). As its value depends on the geo-lithological composition of the growing area, it represents a direct link between a product and its cultivation area. Despite an increase in the scientific literature related to its use as a geographical tracer, little is known about its applicability to timber traceability studies, mostly related to the archaeological domain (English et al., 2001; Erban Kochergina et al., 2021; Hajj et al., 2017). Due to the lack of information from the literature, before conducting our main research study, a preliminary experiment was set up to check the variability of the Sr isotope ratio within a trunk radial section in comparison with the intra-stand variability. To do so, five trunk sections (ca. 30 x 40 cm) of Norway spruce (Picea abies Karst.) belonging to independent trees were collected from a forest stand in South Tyrol, characterized by limestone bedrock. From three sections out of five, a grid sampling (ca. 5 x 5 cm) over the trunk surface was performed by collecting more than 20 independent wood cores randomly selected using a 10-mm increment borer (Haglöf, Sweden) to a depth of ca. 5 cm (Figure A1, panel A). From the remaining two sections, only three independent replicates were collected, without following a specific sampling scheme. Samples were oven-dried (40 °C for 72 h) and finely pulverized in a zirconia ball mill. For each independent sample, an aliquot (ca. 0.5 g) was acid digested in a microwave digestion system and measured at the ICP-MS to determine the Sr concentration. After the Sr/matrix separation, the Sr isotope ratio was measured at the MC ICP-MS. For the detailed method description, refer to the Material and Method section of the main text.

Results indicate that, within a single tree, a certain degree of variability can be observed (Figure A1, panels B:D). The sampling grid did not exactly follow the tree ring distribution in the radial section; nonetheless, it seems that the major variability of the $^{87}Sr/^{86}Sr$ ratio goes from the external growth rings to the pith (transverse surface), while the values seem to remain more constant along the radial section. The average variability along the transverse surface, expressed as the difference between the maximum and minimum measured $^{87}Sr/^{86}Sr$ ratio, is indeed ca. 2-3 times higher compared to the average variability along the radial surface (Figure A1,

Panels B:D). This could be potentially due to the tree growth contemporary to the extension of the root apparatus. We speculate that this temporal trend might be associated with the development of the root apparatus reaching soil layers characterized by different pools of bioavailable Sr. This agrees with the temporal variability observed for the multi-element composition and reported in the literature by other authors (Boeschoten et al., 2023; Rice et al., 2022). To verify this hypothesis, soil samples collected at different depths close to the tree would be needed but this was outside the purpose of the present investigation. In any case, the overall intra-tree variability of the $^{87}Sr/^{86}Sr$ ratio does not exceed the range covered by the values measured in the five available trunk sections. The maximum delta value measured intra-tree was ca. half of the intra-stand delta value (<0.0005 and >0.0009, respectively) (Figure A2). In this forest stand, the $^{87}Sr/^{86}Sr$ ratio measured in the wood samples falls in a rather narrow range, between 0.7089 and 0.7098 (Figure A2), typical for soils developed on this type of substrate (Willmes et al., 2018). The overall low variability measured at this location can be attributed to the bedrock type (limestone), which is generally rather homogeneous. Even though different trends could be observed according to the different bedrock compositions present in mountain areas, we assumed that the intra-tree variability should be always lower or similar to the intra-stand variability, as verified in a study conducted in apple orchards (Aguzzoni et al., 2019). For these reasons, for our main investigation, we drafted a sampling protocol that foresees the sampling of a single wood core from each tree, collected at breast height using an increment borer. From each sample, it was established to remove the bark and further process the whole wood core, including all the tree rings to the pith.

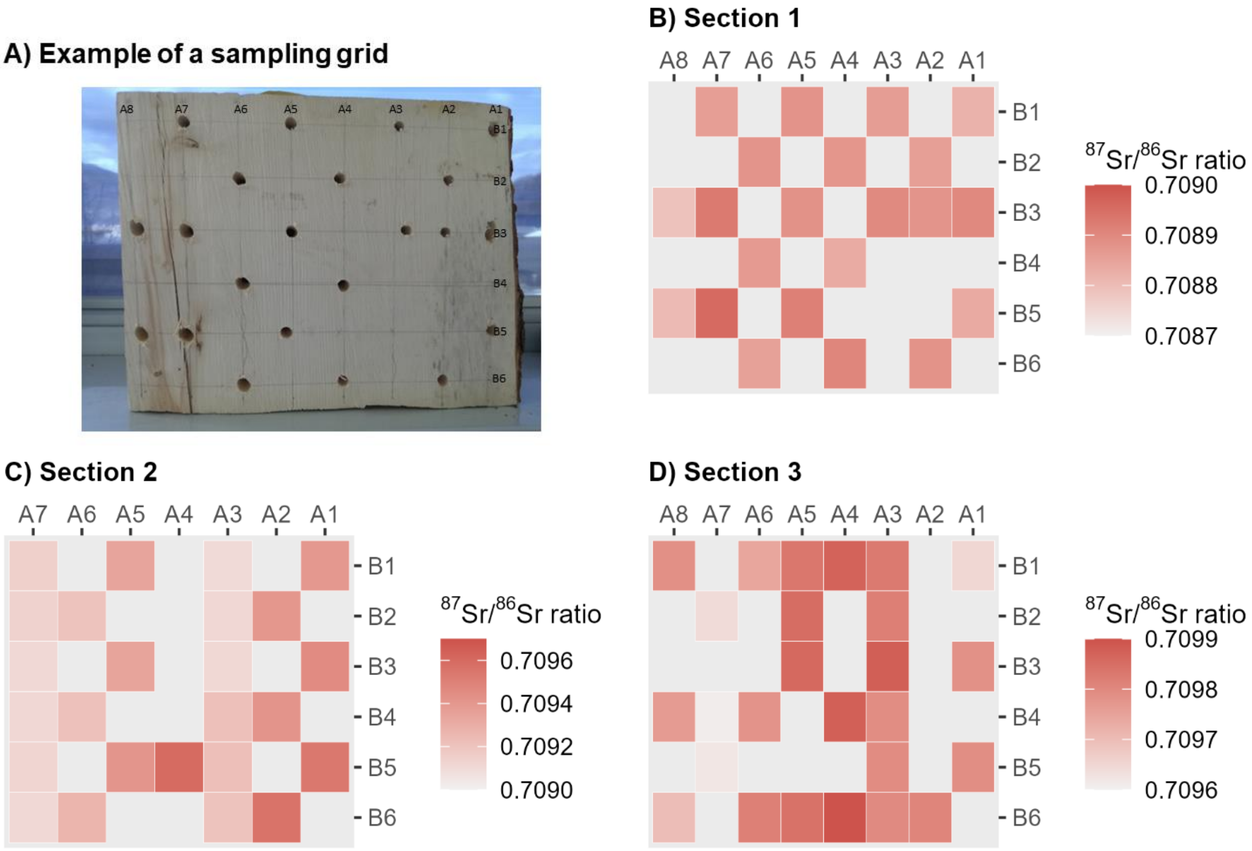


**Figure S1.** Example of a sampling grid distributed over the trunk radial section (Panel A) and results of the Sr isotope ratio analysis for the different sampling points for the three different trunk sections (Panel B:D).

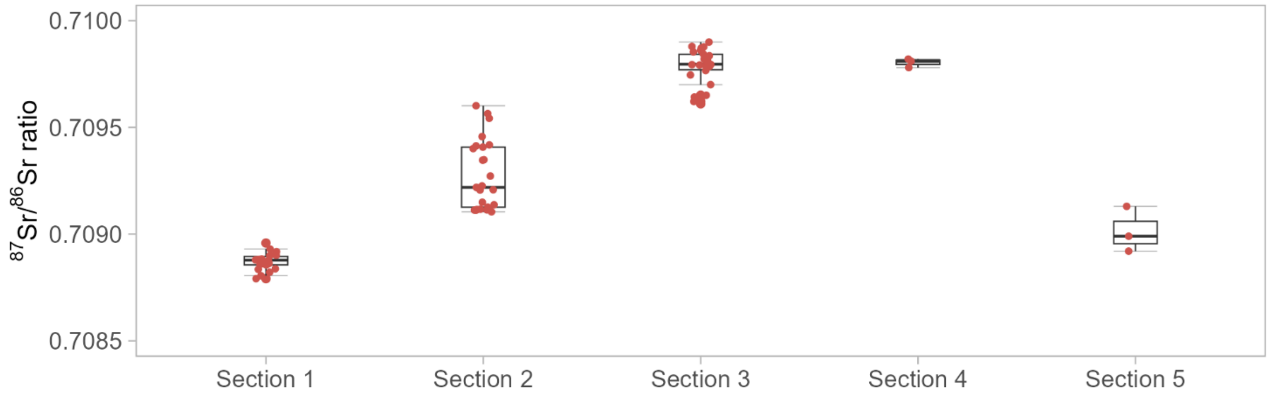


**Figure S2.** Results of the Sr isotope ratio analysis measured in the trunk sections collected for the preliminary experiment (forest stand in South Tyrol, characterized by a limestone bedrock).
